# Low-latitude environmental regularity sustains non-photic entrainment in blind adults

**DOI:** 10.64898/2026.03.19.712663

**Authors:** Karen C. Pugliane, Lucas G. S. França, Mario Leocadio-Miguel, John F. Araújo

## Abstract

The light–dark cycle shaped by Earth’s rotation provided the evolutionary conditions under which circadian rhythms emerged. Consistent with this, previous studies indicate that less than 40% of total blind individuals, who lack photic input, entrain to the 24-h cycle, further evidencing the critical role of light as the dominant zeitgeber for circadian alignment. However, this assumption has been tested almost exclusively in temperate, high-latitude regions, where environmental cues vary seasonally. Near the equator, by contrast, photoperiod and temperature cycles remain exceptionally stable. This highlights a fundamental gap: can circadian rhythms in humans remain synchronised without light when environmental temporal cues are highly regular? We addressed this question in 58 blind adults (21–77 years; 43.1% female) living near the equator in Rio Grande do Norte, Brazil (∼5°S), who wore wrist actigraphy continuously for four weeks. Light sensitivity was assessed through the pupillary light reflex (PLR; 22 PLR-reactive, 36 non-reactive). Applying a semi-supervised machine learning approach to uncover multidimensional patterns without prior categorisation, we identified two distinct phenotypes: a Higher Circadian Stability (HCS; 72%, n = 42) and a Lower Circadian Stability group (LCS; 28%, n = 16). Notably, 64% of PLR-non-reactive individuals (23 of 36) were classified within the HCS group, a proportion approximately 1.6 times higher than previously reported for blind cohorts. These findings demonstrate that, under exceptionally regular equatorial conditions, non-photic cues can sustain a robust circadian entrainment even in the absence of photic input. We propose that environmental regularity promotes the synergy of non-photic timing signals, underscoring ecological context as a key determinant of human circadian temporal organisation.

## Introduction

In humans, biological rhythms align with the light–dark cycle through photic signals detected by the retina and transmitted via the retinohypothalamic tract to the suprachiasmatic nucleus (SCN), which acts as the master pacemaker of the circadian system, coordinating oscillators distributed throughout the body (Skene & Arendt, 2006). When photic input is reduced or absent, circadian entrainment may be compromised, leading to disruptions in physiological and behavioural rhythms.

Studies involving blind individuals frequently report alterations in circadian organisation, typically manifesting free-running patterns and circadian rhythm sleep–wake disorders, such as non-24-hour sleep–wake disorder (Sack et al., 1992; Lockley et al., 1997; Sack et al., 2000; Flynn-Evans & Lockley, 2016). Accordingly, blindness has been conceptualised not only as the loss of vision but also as a condition that may limit access to photic Zeitgeber, essential for maintaining the internal temporal order, thereby increasing physiological vulnerability to circadian misalignment and instability of daily rhythms (Allen, 2019).

Importantly, blindness encompasses a spectrum of visual capacities, ranging from residual light awareness to complete absence of light perception. Although all forms involve impaired image-forming vision (Lee & Mesfin, 2020), the integrity of non-image-forming photoreception may vary substantially. While conscious vision is mediated by outer retinal photoreceptors (rods and cones), non-image-forming responses, such as melatonin suppression and the pupillary light reflex, are essentially driven by intrinsically photosensitive retinal ganglion cells (ipRGCs) located in the inner retina (Feigl & Zele, 2014). When ipRGC pathways are preserved, circadian photoreception can remain functional despite the absence of visual awareness (Czeisler, 1995; Van Gelder, 2008). Evidence from the largest cohort study to date, involving 127 blind women, suggests that circadian photoreception depends not only on the presence of light perception but also on the aetiology of blindness. Damage affecting retinal ganglion cells tends to disrupt circadian alignment, whereas damage restricted to the outer retina may preserve it (Flynn-Evans et al., 2014).

Despite this consensus, data from our laboratory challenged the prevailing view that blindness, particularly in the absence of light perception and responsiveness, almost always disrupts the circadian system. In a pilot study conducted in Natal, Brazil (∼5° S and ∼ 35° W), it was observed that 13 out of 14 totally blind participants without pupillary reactivity exhibited 24-hour period of rest-activity rhythm (RAR), with activity properly aligned to daytime hours (Depieri, 2021). We suggest that this finding should be interpreted in light of the geographical context in which the study was conducted. Most studies reporting a high prevalence of free-running rhythms among blind populations have been performed at higher latitudes, particularly in North America and Europe (e.g., Sack et al., 1992; Klerman et al., 1998; Skene et al., 1999; Léger et al., 2002; Emens et al., 2013; Flynn-Evans et al., 2014), where pronounced seasonal variation in photoperiod and temperature may impose additional challenges on the circadian timing system, potentially reducing the stability of entrainment.

Key circadian parameters, including melatonin secretion and nocturnal rest duration, are modulated by daylength and seasonal changes, which can influence the phase and organisation of endogenous rhythms (Cipolla-Neto & Amaral, 2018; Monsivais-Velazquez et al., 2017). Population-based studies further indicate that geographic context and latitude are associated with measurable differences in sleep behaviour and circadian outcomes, supporting the role of environmental factors in shaping human temporal organisation (DelRosso & Vodapally, 2026) In contrast, equatorial regions, where our study was conducted, exhibit minimal seasonal variation in photoperiod, remaining close to 12:12 throughout the year (Supplementary Figure 1). Furthermore, solar radiation and ambient temperature display pronounced daily oscillations, generating stable and predictable environmental signals that may contribute to circadian entrainment.

We previously proposed (Leocadio-Miguel et al., 2017) that proximity to the Equator may favour stronger circadian entrainment due to both phasic (light transitions) and tonic (irradiance intensity) influences of light, which in turn results in higher prevalence of morning chronotypes (Randler, 2008; Randler et.al., 2017). As mean solar irradiance decreases with latitude, the coupling between circadian oscillators may be altered; leading to slower oscillator velocity, longer circadian periods, and a delayed-phase pattern compared to low-latitude conditions. This phenomenon suggests that environmental consistency may facilitate rhythm consolidation (Díez-Noguera, 1994; Hut et.al., 2013). However, whether environmental regularity can compensate for reduced photic input in blind individuals, remains largely unknown.

To expand the current evidence and address these discrepancies, we characterised circadian rest–activity rhythms (RAR) in people with blindness residing near the Equator. Actigraphy combined with non-parametric metrics was used to preserve essential features of the RAR, including its asymmetry and irregular transitions between rest and activity phases, which are not well captured by sinusoidal models (Witting et al., 1990; Van Someren et al., 1999; Gonçalves et al., 2015). Pupillary light reactivity (PLR) was employed as a physiological proxy of melanopsin photoreception (Markwell et al., 2010; Zaidi et al., 2007; Bonmati-Carrion et al., 2016), reflecting the functional status of ipRGC signalling.

We hypothesised that: (i) a refined, data-driven analytical framework would detect subtle alterations in circadian rhythmicity overlooked by parametric approaches, and (ii) multivariate rhythmic profiles would reveal distinct circadian phenotypes that extend beyond PLR status. To test these hypotheses, we explored a comprehensive suite of metrics, including stability, fragmentation, amplitude, and phase markers (e.g., L5/M10 onset and sleep midpoint), alongside light exposure and frequency domain parameters. Finally, we applied a semi-supervised machine learning approach to identify latent circadian profiles, revealing phenotypes characterised by distinct patterns in rest–activity features.

## Methods

### Geographic and Climatic Parameters

This study was conducted in Natal, a coastal tropical city in northeastern Brazil. Data were collected between April–November 2023 and April–September 2024. Participants resided in both Natal and surrounding municipalities within the state of Rio Grande do Norte (Supplementary Figure S2). All analyses were conducted in Python (v3.12.12). Geospatial processing was performed using geopandas (v1.1.0) and shapely (v2.1.0). Official spatial datasets from the Brazilian Institute of Geography and Statistics (IBGE) were accessed using the geobr package. The centroid of each municipality polygon was used as the geographic reference for environmental and solar calculations. Across the study area, municipality centroids ranged from 5.41°S to 6.36°S in latitude and 35.23°W to 36.82°W in longitude.

Climatic conditions were assessed using daily meteorological data from the NASA POWER Project API (https://power.larc.nasa.gov), while sunrise and sunset times were calculated using the Astral library (v3.2;https://astral.readthedocs.io) based on municipality centroid coordinates. During the collection period, the average sunrise time was 05:20 ± 11 min (range 04:54–05:32) and the average sunset time 17:17 ± 3 min (range 17:12–17:22), yielding an approximately 12 h light–dark cycle (mean photoperiod 12.11 ± 0.23 h, range 11.75–12.48 h), corresponding to an annual photoperiod amplitude of approximately 44 minutes. Across municipalities and months, mean air temperature averaged 27.01 ± 1.07 °C (range 24.42–30.36 °C), while mean surface solar radiation averaged 243.59 ± 24.65 W/m² (range 187.73–296.11 W/m²) confirming the low seasonal environmental variability typical of tropical low-latitude environments (Supplementary Figure S1).

### Participants

Data collection was carried out at the Institute of Education and Rehabilitation of the Blind of Natal (IERC), a non-profit institution for individuals with visual impairment. A total of 64 volunteers aged between 21 and 77 years were invited to participate in the study. All volunteers were recruited from the institution’s official registry, which includes individuals with medically diagnosed blindness or legal blindness, evaluated by qualified healthcare professionals. The study was approved by the Human Research Ethics Committee of the Federal University of Rio Grande do Norte (protocol number CAAE: 60610322.7.0000.5537) and conducted in accordance with the Declaration of Helsinki. Brazilian ethical regulations for human research (Resolution No. 466/2012, National Health Council) were followed, and all participants were voluntarily enrolled without financial compensation or inducement. Informed consent was obtained through written and verbal procedures; for participants unable to provide a signature, consent was confirmed using fingerprint identification.

### Assessment of Pupillary Light Reflex (PLR)

Pupillary reactivity was assessed based on pupillary constriction in response to a flashlight. The flashlight was battery-powered and equipped with a white LED light source (600 lumens). Assessments were performed in a darkened room to minimise ambient light. The examiner directed the flashlight toward one eye at approximately 10 cm, and pupillary constriction was visually inspected. Participants were classified as reactive when a clear constriction was observed in response to the light stimulus, and as non-reactive when no constriction was detected (Markwell et al., 2010; Smith et al., 2020).

Participants were also interviewed regarding their conscious light perception, specifically whether they could perceive light or distinguish between light and dark environments. This self-reported information was recorded and used for supplementary analyses. Visual acuity was further assessed using a Snellen chart in combination with structured questionnaire items adapted from standard clinical assessment guidelines (MSD Manual Professional Edition: Assessing Visual Acuity). Among the 58 participants included in the sample, twenty nine were unable to perceive light, twenty three retained minimal visual function (light perception, hand motion perception, or counting fingers), and six were able to identify the largest optotype.

For analytical purposes, pupillary light reactivity (PLR) was used as a proxy for intrinsically photosensitive retinal ganglion cell (ipRGC) integrity and served as the primary grouping variable in subsequent analyses.

### Actigraphy Protocol and Data Processing

Participants were invited to wear an actigraphy device (ActTrust 1; Condor Instruments, São Paulo, Brazil) continuously for four consecutive weeks, placed on the non-dominant wrist, defined as the arm not used to handle the mobility cane. They were instructed to keep the monitor on throughout the entire day, removing it only during bathing or high-impact physical activities. A verbal description of the device and its functioning was provided before the start of the monitoring period. Weekly in-person follow-up meetings were conducted at the Institute for the Education and Rehabilitation of the Blind to verify proper actigraphy signal acquisition.

For data processing, participants were excluded if they did not have valid actigraphy data for at least 10 days (total of 6 exclusions), as recommended for circadian rhythm assessment (Refinetti et.al., 2007). A valid day was defined as one in which the device was worn for at least two-thirds of the time during both the light and dark phases (i.e., ≥16 hours per day; Danilevicz et al., 2024). In addition, periods containing three or more consecutive invalid days were removed from the time series data.

### Actigraphy-Derived Variables

Circadian rest–activity rhythmicity was assessed through non-parametric methods (Gonçalves et al., 2015) and frequency-domain analyses (Sokolove & Bushell, 1978). We used the PyActigraphy Python library (Hammad et al., 2021) to evaluate a set of metrics designed to characterise circadian rhythmicity across multiple physiological signals, all recorded with a wrist-worn actigraphy device equipped with activity, ambient light, and skin temperature sensors; See the Zenodo repository (https://doi.org/10.5281/zenodo.18922404) for full code.

Metrics related to stability, fragmentation, amplitude, and phase timing were derived from rest–activity data. Measures of light exposure were computed from the actigraph’s light sensor, while frequency-domain characteristics—such as rhythmic prominence and estimated period—were assessed across activity, light, and peripheral skin temperature signals (Table 1).

**Table 1.**
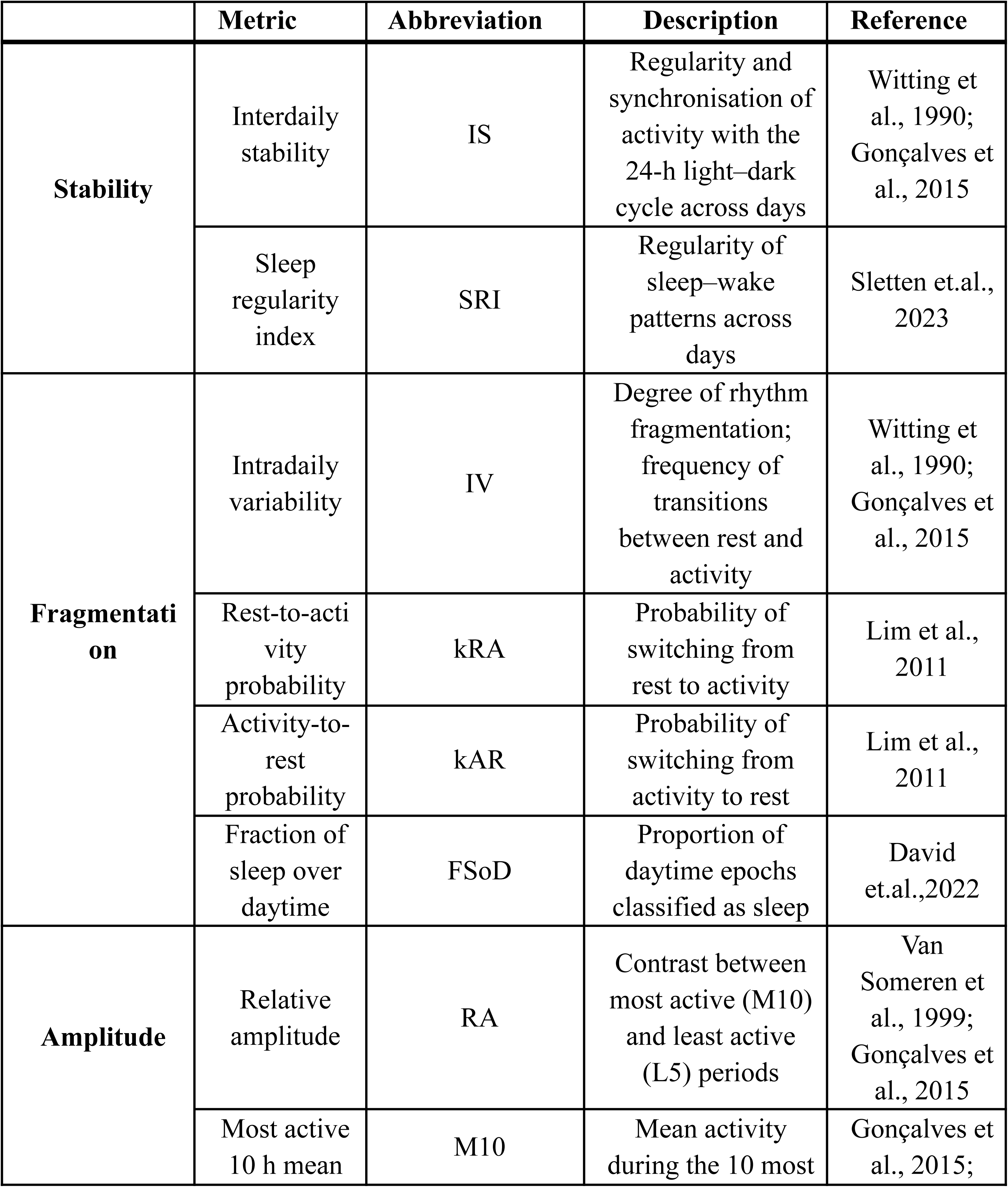
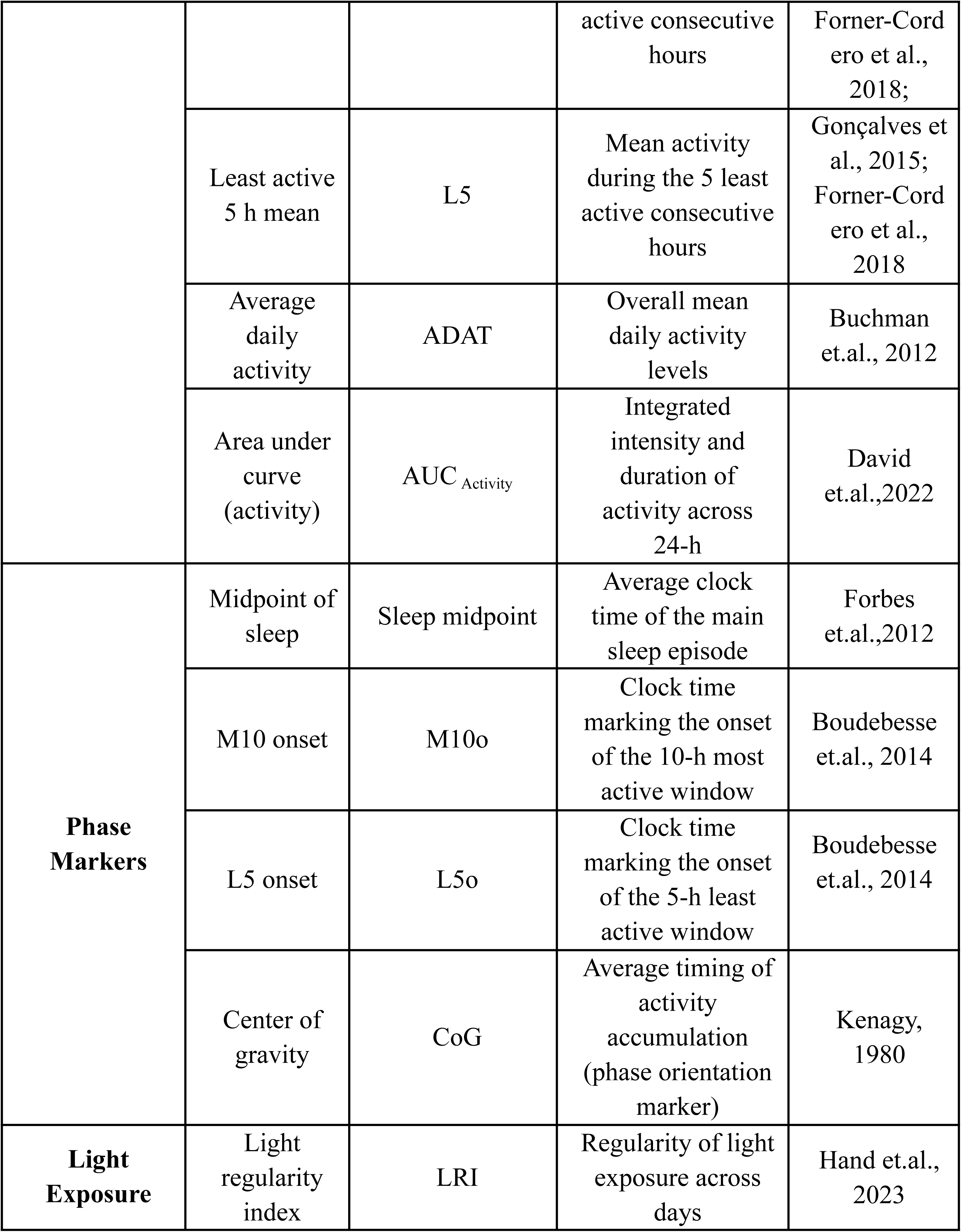
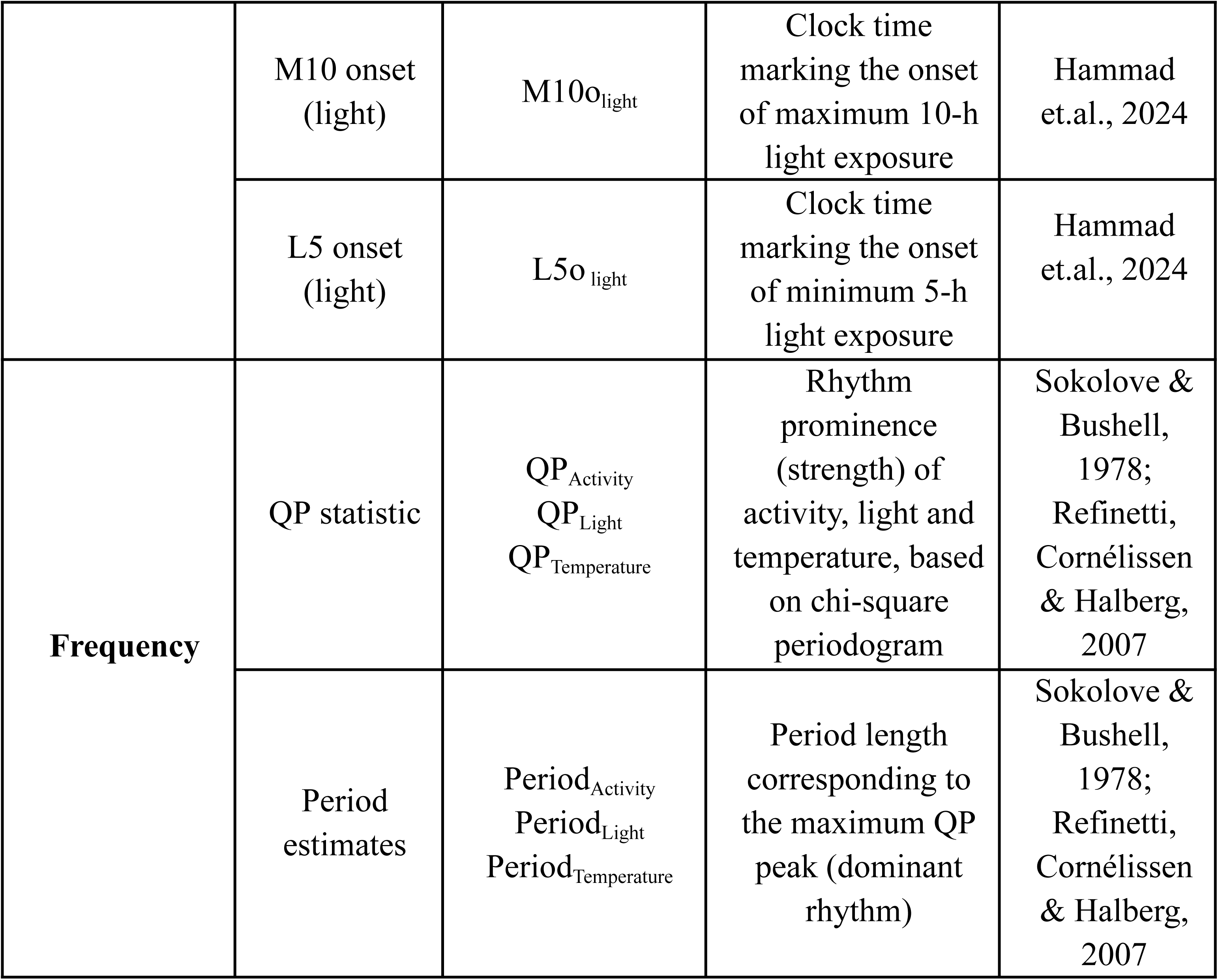
Description of non-parametric and frequency-domain metrics derived from actigraphy.

### Statistical Analysis

All analyses were conducted in Python (v3.12.12) using standard scientific computing libraries, including pandas (v2.2.2), numpy (v2.0.2), scipy (v1.16.3), statsmodels (v0.14.6), scikit-learn (v1.6.1), seaborn (v0.13.2), matplotlib (v3.10.0), and shap (v0.51.0). Descriptive statistics were calculated to assess sample distribution and data integrity. Categorical variables were compared using chi-square tests. For continuous variables, normality was assessed using the Shapiro–Wilk test, and homogeneity of variance was evaluated with Levene’s test.

All 24 variables listed in Table 1 were included in the initial analysis and standardised using Z-score normalisation. Multicollinearity was assessed via Pearson correlation matrices (threshold > 0.85) and Variance Inflation Factor (VIF > 5.0). Variables meeting both criteria were excluded. This resulted in the exclusion of AUC due to redundancy with ADAT. Principal Component Analysis (PCA) was then applied for dimensionality reduction of the 23 remaining variables. The number of components was determined as the minimum required to explain ≥80% of the cumulative variance (Jolliffe & Cadima, 2016).

Upon reducing the dimensionality of our dataset, we deployed an unsupervised machine learning method (K-means) for a data-driven approach. We evaluated *k* values ranging from 2 to 10. The optimal number of clusters was selected based on the highest silhouette score. To assess the contribution of each feature to cluster separation, we employed a semi-supervised machine learning approach. After defining cluster membership, a Random Forest classifier was trained using the cluster labels as targets, allowing the estimation of feature importance using SHAP values (Ponce-Bobadilla et.al., 2024).

Between-cluster comparisons were performed across all selected numeric variables using either two-tailed independent-samples *t*-tests or Mann–Whitney *U* tests, depending on the normality and homogeneity of variance assumptions for each variable. Categorical variables were assessed using chi-squared tests. Within-cluster associations among circadian metrics were explored using pairwise Spearman correlations, with FDR correction applied to control for multiple testing. Finally, linear regression models were constructed to examine whether cluster membership and pupillary reactivity predicted key circadian outcomes.

We then fitted a Gaussian generalised linear model (identity link) for each selected outcome:

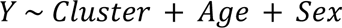

where Cluster was treated as a categorical binary predictor (HCS/LCS), Age was continuous, and Sex was binary (Female/Male). Models were estimated using HC3 heteroscedasticity-consistent (robust) standard errors, reducing sensitivity to heteroscedasticity and mild deviations from classical assumptions. For each outcome, we extracted the cluster coefficient (β; Cluster HCS vs Cluster LCS), its 95% confidence interval, and the corresponding t-statistic and p-value. Full results for this model are provided in the Supplementary Material (Table S2).

All tests were two-tailed, with statistical significance set at α = 0.05. To account for multiple testing across outcomes, all p-values were adjusted for multiple comparisons using the Benjamini–Hochberg False Discovery Rate (FDR) correction (Benjamin & Hochberg, 1995)

The data analysis scripts developed for this study are publicly available at: https://doi.org/10.5281/zenodo.18922404.

## Results

### Sample Characterisation and Circadian Period Classification

The final sample comprised 58 participants, classified as either reactive (with pupillary light reactivity, *n* = 22) or non-reactive (without pupillary light reactivity, *n* = 36), with ages ranging from 21 to 77 years. No significant differences were found between groups in age or sex distribution (all *p* > 0.05; see Table 2). The reactive group included 12 females (54.6%), while the non-reactive group included 13 females (36.1%). Participants wore the actigraphy devices for a period ranging from 10 to 60 days, with a mean of 28 days (*SD* = 7.6). Regarding the circadian period of the activity–rest rhythm, we estimated the most prominent periodicity using the chi-square periodogram, as implemented by Sokolove and Bushell (1978).

**Table 2.**
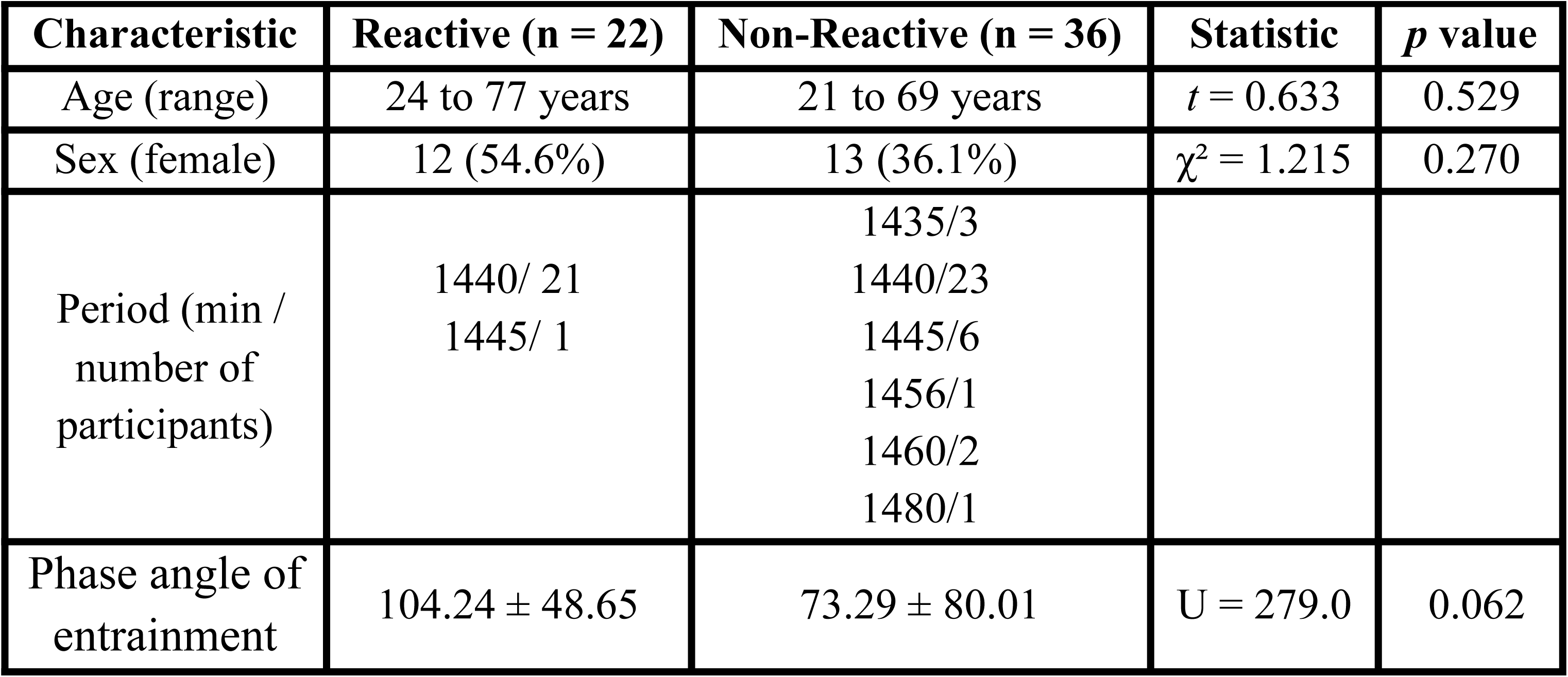
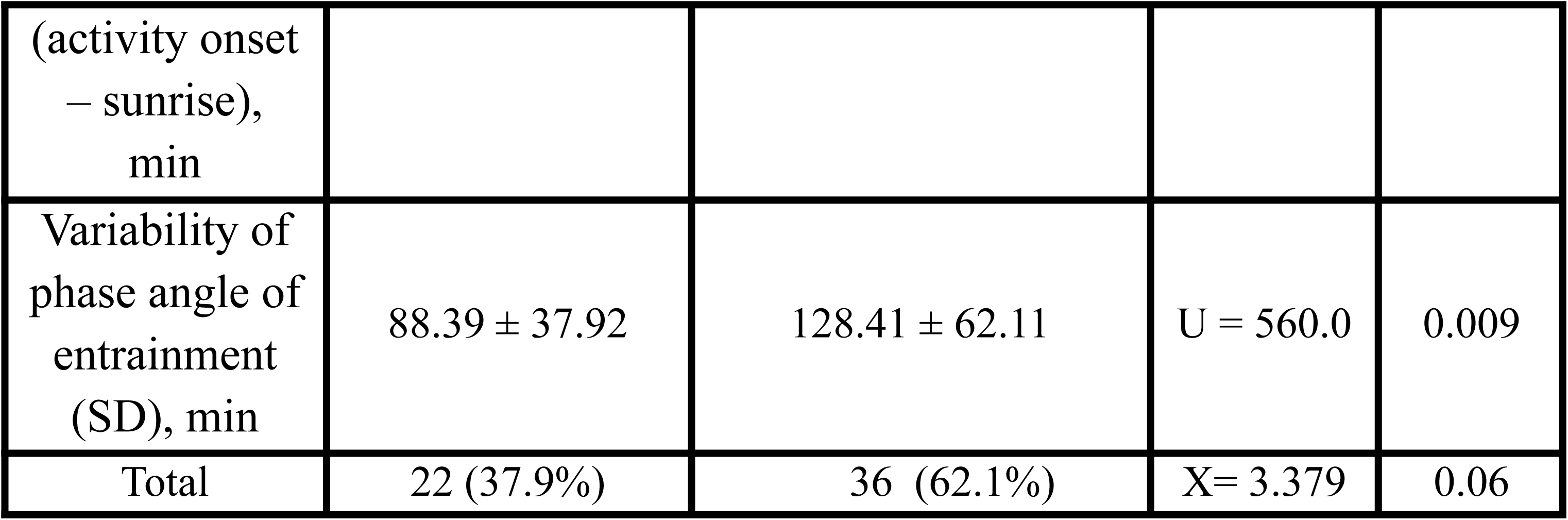
Characteristics of participants according to pupillary light reactivity.

Participants were considered to have a non-entrained rhythm if their circadian period fell outside the range of 23.88–24.12 h, a threshold based on the normal variance of the intrinsic circadian period (𝛕) observed in sighted individuals (Wright et al., 2001). This criterion has been previously applied in blind cohorts using circadian phase markers (such as melatonin) to define entrainment (Flynn-Evans et al., 2014) and was applied here to activity–rest rhythms as a behavioral proxy for circadian organisation. We found that all participants in the reactive group exhibited entrained circadian rhythms. In the non-reactive group, 80.6% were entrained and 19.4% were non-entrained. (Table 2).

To characterise the temporal alignment between behavioral activity and the environmental light–dark cycle, we calculated the phase angle (φ) between activity onset and local sunrise (Figure S3). Across participants, individual mean phase angles ranged from − 98 min to 244.6 min, corresponding to activity onset occurring from approximately 1 h 38 min before sunrise to 4 h 05 min after sunrise (roughly 03:40 to 09:25 local time). Participants without a pupillary light response showed a mean phase angle of 73.3 ± 80.0 min, whereas reactive participants showed a mean phase angle of 104.2 ± 48.7 min. Although reactive participants tended to initiate activity later relative to sunrise, this difference was not statistically significant (Mann–Whitney U = 279, p = 0.062).

Furthermore, differences emerged when examining phase angle stability (SD). The variability of the phase angle across days was significantly greater in the non-reactive group (128.4 ± 62.1 min) compared with the reactive group (88.4 ± 37.9 min; Mann–Whitney U = 560, p = 0.009). Together, these results indicate that while the average alignment of activity onset with sunrise did not differ significantly between PLR groups, participants lacking a pupillary light response exhibited greater day-to-day variability in the temporal alignment between behavior and the solar cycle.

### Distinct Circadian Profiles Identified Through Cluster Analysis

A total of 23 variables extracted from the actigraphy data was retained for analysis after multicollinearity screening. With PCA, we considered the first 9 components, explaining 80.02% of the total variance. Furthermore, our cluster analysis suggested an optimal number of clusters (*k* = 2) with a silhouette score = 0.2452 (Supplementary Figure S4). Two distinct circadian profiles emerged through unsupervised clustering based on multivariate rhythm metrics, as visualised in the PCA space (Figure 1A). Group labels were subsequently defined based on feature importance.

**Figure 1.**
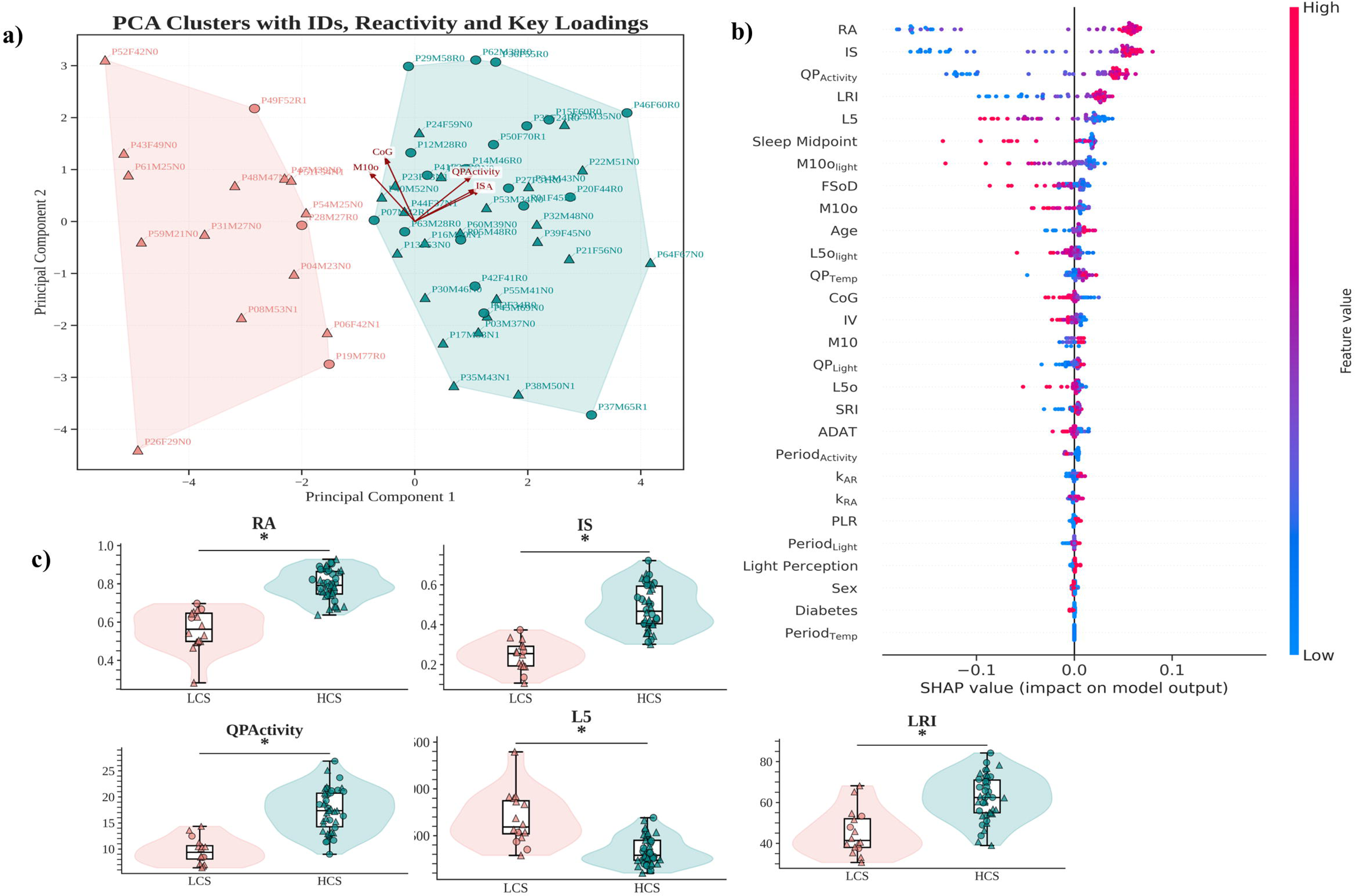
Circadian Cluster Characterisation. **(a)** PCA plot illustrating the separation between the two identified clusters (k = 2) based on rest-activity rhythm variables. Each point represents an individual participant. Colors denote cluster membership (HCS: Higher Circadian Stability; LCS: Lower Circadian Stability), and shapes indicate pupillary reactivity (circles: reactive; triangles: non-reactive). Arrows represent the loadings of the five most influential variables in the PCA space (QPActivity, RA, Ml Oo, CoG, and IS), with direction and length indicating their contribution to the principal components. **(b)** SHAP beeswarm plot showing the contribution of each variable to the supervised model used to distinguish the clusters identified by the unsupervised clustering procedure. **(c)** Violin-box plots comparing the variables with the highest SHAP importance between clusters of circadian stability. The violin plots represent the distribution of values, boxplots indicate the median and interquartile range, and points represent individual participants. Shapes indicate pupillary reactivity (circles: reactive; triangles: non-reactive).

A Random Forest classifier was trained using the cluster labels as targets, and SHAP values were computed to estimate the relative importance of each feature in distinguishing between clusters. The top three variables ranked by mean absolute SHAP values were RA (0.163), IS (0.143), and QP_Activity_ (0.109). Additional contributors included LRI (0.067) and L5 (0.064) (Figure 2). Based on the predominance of features related to rhythm regularity and amplitude among the top-ranked variables, the two clusters were labelled Higher Circadian Stability (HCS) and Lower Circadian Stability (LCS) (Figure 1B).

**Figure 2.**
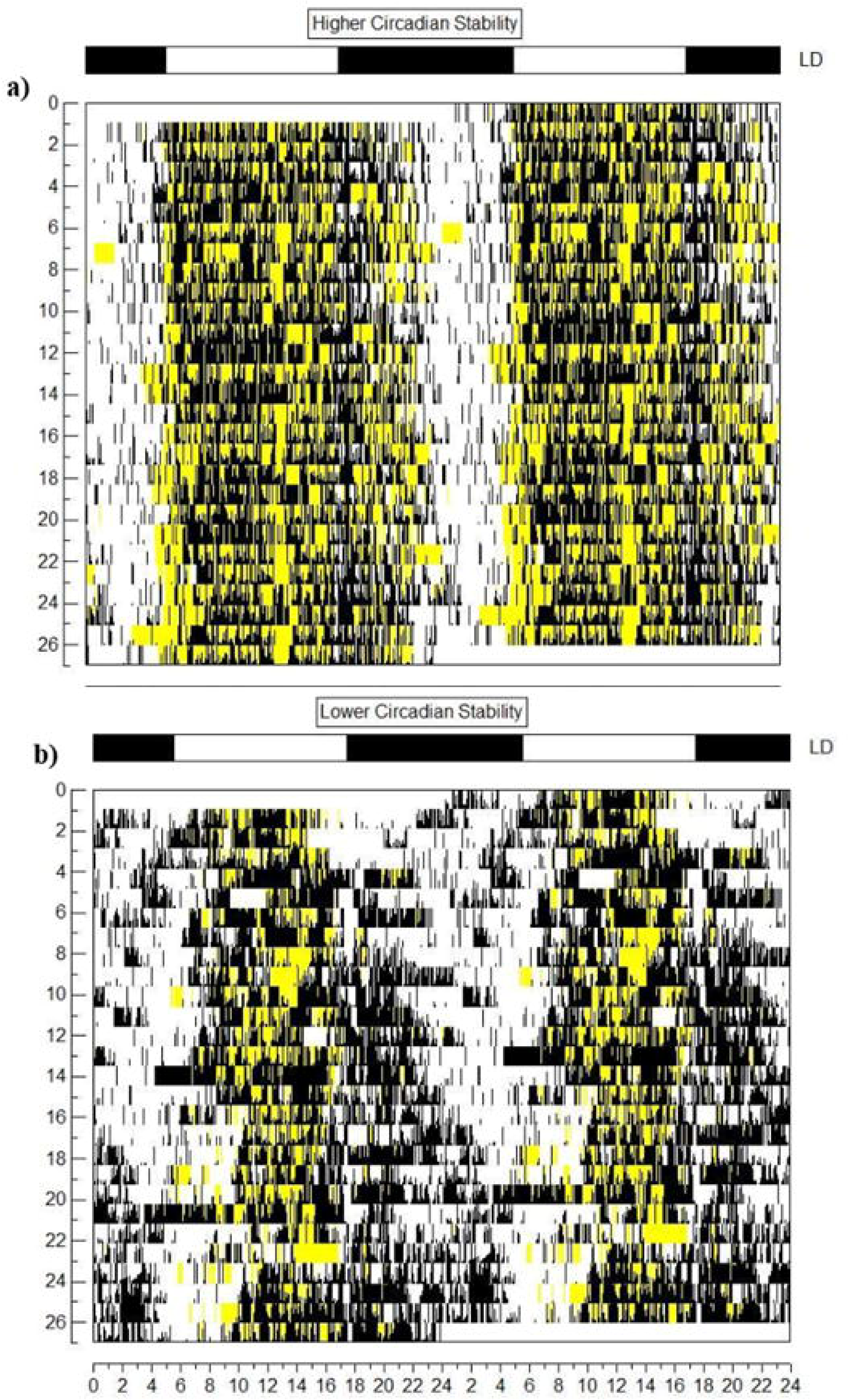
Representative double-plotted actograms from pa1ticipants classified as higher circadian stability (HCS; a) and lower circadian stability (LCS; b). Each row con-esponds to two consecutive days of wrist actigraphy data, with activity counts plotted against time of day (0-24 h). Black bars represent periods of activity, and white areas represent rest. Light exposure of the pa1ticipant is highlighted in yellow. Pa1ticipant P3 (bilateral enucleation, no pupilla1y light reflex) is shown as an example of the HCS group, a pa1ticipant P31 (one enucleated eye, no pupilla1y light reflex) illustrates the LCS group.

### Comparison Between Circadian Clusters / Characterisation of Circadian Clusters

Among the 23 actigraphy-derived variables evaluated, 14 showed statistically significant differences between clusters. In circadian stability measures, the HCS shows higher Interdaily Stability and (t = 13.00, p < 0.001) and Sleep Regularity Index (t = –2.74, p = 0.008). We also have higher amplitude (t = 9.38, p < 0.001) and lower rest activity L5 (t = –594.00, p < 0.001). In terms of rhythm fragmentation, the HCS group also exhibited lower FSoD values (t = –490.00, p = 0.008), suggesting more consolidated rest-activity rhythms.

For light exposure variables, the HCS group showed more regular light exposure patterns (LRI: t = 5.38, p < 0.001), earlier exposure onset (M10o: t = –4.23, p < 0.001), and earlier exposure offset (L5olight: t = –2.77, p = 0.008). Regarding activity phase markers, the HCS group also demonstrated earlier timing of activity onset (M10o: t = –4.23, p < 0.001). In the frequency domain, significant differences were observed in PeriodActivity (t = –467.00, p = 0.002), QPActivity (t = 30.00, p < 0.001), and QPTemperature (t = 2.88, p = 0.006), further differentiating the clusters. Furthermore, two additional variables: ADAT (t = –2.24, p = 0.029) and CoG (t = –2.04, p = 0.046), showed between-group differences but did not survive FDR correction. These patterns reinforce the interpretation that the clustering solution captured individuals with stable versus fragmented circadian profiles (Table 3).

**Table 3:**
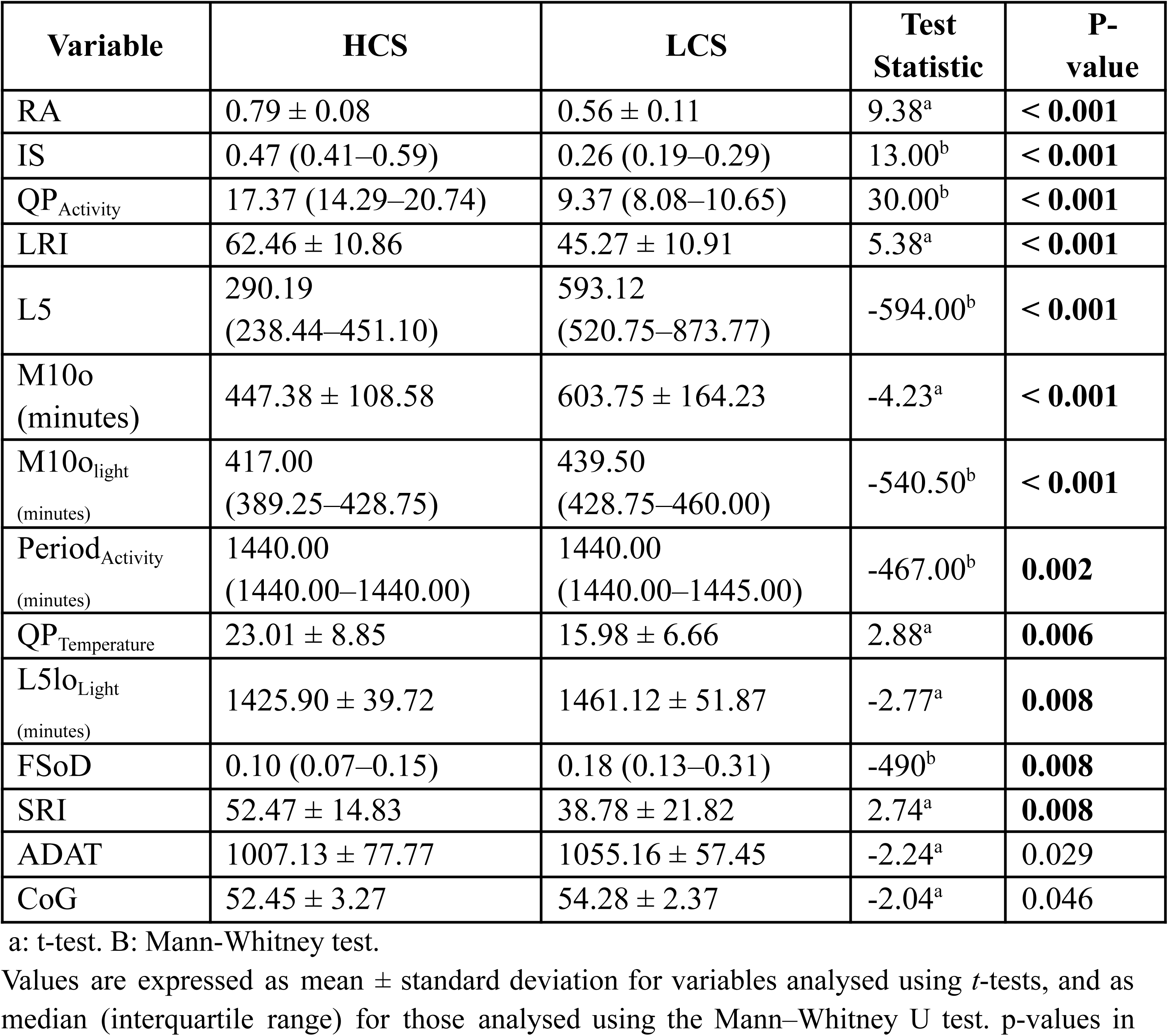
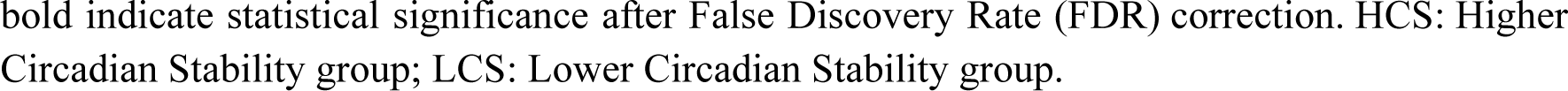
Comparison of circadian rhythm metrics between higher and lower circadian stability groups (HCS vs. LCS)

| Variable | HCS | LCS | Test Statistic | P-value |
| --- | --- | --- | --- | --- |
| RA | 0.79 ± 0.08 | 0.56 ± 0.11 | 9.38<sup>a</sup> | < **0.001** |
| IS | 0.47 (0.41–0.59) | 0.26 (0.19–0.29) | 13.00<sup>b</sup> | < **0.001** |
| QP<sub>Activity</sub> | 17.37 (14.29–20.74) | 9.37 (8.08–10.65) | 30.00<sup>b</sup> | < **0.001** |
| LRI | 62.46 ± 10.86 | 45.27 ± 10.91 | 5.38<sup>a</sup> | < **0.001** |
| L5 | 290.19<br>(238.44–451.10) | 593.12<br>(520.75–873.77) | -594.00<sup>b</sup> | < **0.001** |
| M10o (minutes) | 447.38 ± 108.58 | 603.75 ± 164.23 | -4.23<sup>a</sup> | < **0.001** |
| M10o<sub>light</sub> (minutes) | 417.00<br>(389.25–428.75) | 439.50<br>(428.75–460.00) | -540.50<sup>b</sup> | < **0.001** |
| Period<sub>Activity</sub> (minutes) | 1440.00<br>(1440.00–1440.00) | 1440.00<br>(1440.00–1445.00) | -467.00<sup>b</sup> | **0.002** |
| QP<sub>Temperature</sub> | 23.01 ± 8.85 | 15.98 ± 6.66 | 2.88<sup>a</sup> | **0.006** |
| L5lo<sub>Light</sub> (minutes) | 1425.90 ± 39.72 | 1461.12 ± 51.87 | -2.77<sup>a</sup> | **0.008** |
| FSoD | 0.10 (0.07–0.15) | 0.18 (0.13–0.31) | -490<sup>b</sup> | **0.008** |
| SRI | 52.47 ± 14.83 | 38.78 ± 21.82 | 2.74<sup>a</sup> | **0.008** |
| ADAT | 1007.13 ± 77.77 | 1055.16 ± 57.45 | -2.24<sup>a</sup> | 0.029 |
| CoG | 52.45 ± 3.27 | 54.28 ± 2.37 | -2.04<sup>a</sup> | 0.046 |

a: t-test. B: Mann-Whitney test.
Values are expressed as mean ± standard deviation for variables analysed using *t*-tests, and as median (interquartile range) for those analysed using the Mann-Whitney U test. p-values in
bold indicate statistical significance after False Discovery Rate (FDR) correction. HCS: Higher Circadian Stability group; LCS: Lower Circadian Stability group.

**Table 4.**
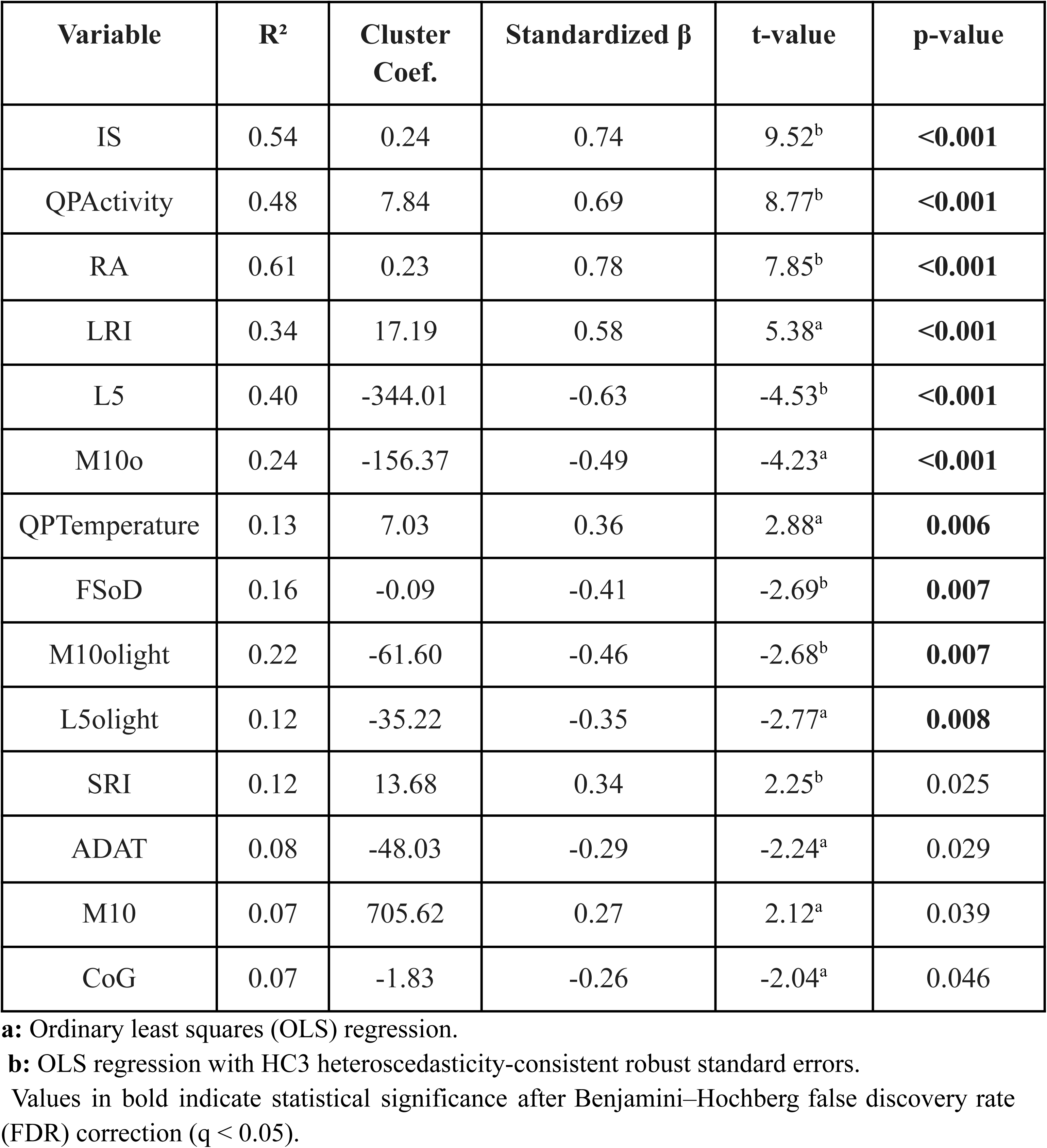
Regression Results Using Cluster Membership as Predictor.

| Variable | R<sup>2</sup> | Cluster Coef. | Standardized $\beta$ | t-value | p-value |
| --- | --- | --- | --- | --- | --- |
| IS | 0.54 | 0.24 | 0.74 | 9.52<sup>b</sup> | **<0.001** |
| QPActivity | 0.48 | 7.84 | 0.69 | 8.77<sup>b</sup> | **<0.001** |
| RA | 0.61 | 0.23 | 0.78 | 7.85<sup>b</sup> | **<0.001** |
| LRI | 0.34 | 17.19 | 0.58 | 5.38<sup>a</sup> | **<0.001** |
| L5 | 0.40 | -344.01 | -0.63 | -4.53<sup>b</sup> | **<0.001** |
| M10o | 0.24 | -156.37 | -0.49 | -4.23<sup>a</sup> | **<0.001** |
| QPTemperature | 0.13 | 7.03 | 0.36 | 2.88<sup>a</sup> | **0.006** |
| FSoD | 0.16 | -0.09 | -0.41 | -2.69<sup>b</sup> | **0.007** |
| M10olight | 0.22 | -61.60 | -0.46 | -2.68<sup>b</sup> | **0.007** |
| L5olight | 0.12 | -35.22 | -0.35 | -2.77<sup>a</sup> | **0.008** |
| SRI | 0.12 | 13.68 | 0.34 | 2.25<sup>b</sup> | 0.025 |
| ADAT | 0.08 | -48.03 | -0.29 | -2.24<sup>a</sup> | 0.029 |
| M10 | 0.07 | 705.62 | 0.27 | 2.12<sup>a</sup> | 0.039 |
| CoG | 0.07 | -1.83 | -0.26 | -2.04<sup>a</sup> | 0.046 |

**a:** Ordinary least squares (OLS) regression.
**b:** OLS regression with HC3 heteroscedasticity-consistent robust standard errors.
Values in bold indicate statistical significance after Benjamini–Hochberg false discovery rate (FDR) correction ( $q < 0.05$ ).

**Table 5.** Regression Results Using Pupillary Reactivity as Predictor.

| **Variab**<br>**le** | **R<sup>2</sup>** | **Coef. PLR** | **Standardize**<br>**d $\beta$** | **t**<br>**(regression)** | **p**<br>**(regressio**<br>**n)** |
| --- | --- | --- | --- | --- | --- |
| RA | 0.15 | 0.11 | 0.39 | **3.59<sup>b</sup>** | **<0.001** |
| L5 | 0.11 | -169.94 | -0.34 | -3.05<sup>b</sup> | **0.002** |
| FSoD | 0.11 | -0.06 | -0.33 | -2.71<sup>b</sup> | 0.007 |
| IS | 0.12 | 0.10 | 0.34 | 2.70<sup>a</sup> | 0.009 |
| LRI | 0.10 | 8.48 | 0.31 | 2.46<sup>a</sup> | 0.017 |
| SRI | 0.10 | 11.40 | 0.31 | 2.45<sup>a</sup> | 0.017 |

**a:** Ordinary least squares (OLS) regression.
**b:** OLS regression with HC3 heteroscedasticity-consistent robust standard errors.
Values in bold indicate statistical significance after Benjamini–Hochberg false discovery rate (FDR) correction ( $q < 0.05$ ).

In this context, group differences were observed across all five variables that most contributed to cluster separation (Figure 1C), as indicated by the SHAP values (Supplementary Table S1). These differences are further illustrated by representative actograms from participants belonging to the HCS and LCS groups (Figure 2). Notably, both participants were non-reactive, and the volunteer in the HCS group presented bilateral enucleation.

### Internal Organisation

Intra-cluster correlation analyses were performed to examine the internal organisation of circadian variables within each group. In the HCS cluster, multiple significant correlations were identified across variables related to stability, amplitude, fragmentation, phase markers, light exposure, and frequency-domain metrics (FDR-corrected *p* < 0.05; Figure 5).

Among stability features, higher interdaily stability (IS) was positively associated with rhythm prominence of activity/rest, relative amplitude, and mean activity during the 10 most active consecutive hours, while showing a negative association with intradaily variability (IS–QPActivity ρ = 0.66; IS–RA ρ = 0.46; IS–M10 ρ = 0.44; IS–IV ρ = –0.45). Similarly, greater sleep regularity, measured by the Sleep Regularity Index (SRI), was associated with a lower probability of transitions between behavioral states, while the fraction of sleep occurring during daytime (FSoD) was negatively associated with the light exposure regularity index (LRI) (SRI–kAR ρ = –0.55; FSoD–LRI ρ = –0.49).

Within amplitude metrics, greater relative amplitude (RA) was associated with higher activity during the most active hours and stronger rhythm prominence, while showing negative associations with activity during the least active 5-h period and behavioral fragmentation (RA–M10 ρ = 0.43; RA–QPActivity ρ = 0.47; RA–L5 ρ = –0.60; RA–kAR ρ = –0.48). Likewise, M10 was positively associated with rhythm prominence and negatively associated with fragmentation (M10–QPActivity ρ = 0.59; M10–IV ρ = –0.52). In contrast, L5 was positively correlated with average daily activity (ADAT), which in turn was associated with greater behavioral state transitions (L5–ADAT ρ = 0.47; ADAT–kAR ρ = 0.44).

Regarding phase markers, the center of gravity of activity (CoG) showed positive associations with both behavioral and light exposure timing markers, indicating coherence between activity timing and environmental light exposure (M10olight–CoG ρ = 0.56; L5olight–CoG ρ = 0.45; SleepMidpoint–CoG ρ = 0.75; L5o–CoG ρ = 0.50). This alignment was further supported by correlations between light exposure phase markers and behavioral phase markers (M10olight–M10o ρ = 0.58; M10olight–SleepMidpoint ρ = 0.43; L5olight–L5o ρ = 0.67; L5olight–SleepMidpoint ρ = 0.42).

Finally, among frequency-domain variables, PeriodActivity was positively correlated with both PeriodLight and PeriodTemperature (PeriodActivity–PeriodLight ρ = 0.42; PeriodActivity–PeriodTemperature ρ = 0.45) (Figure 3A).

**Figure 3.**
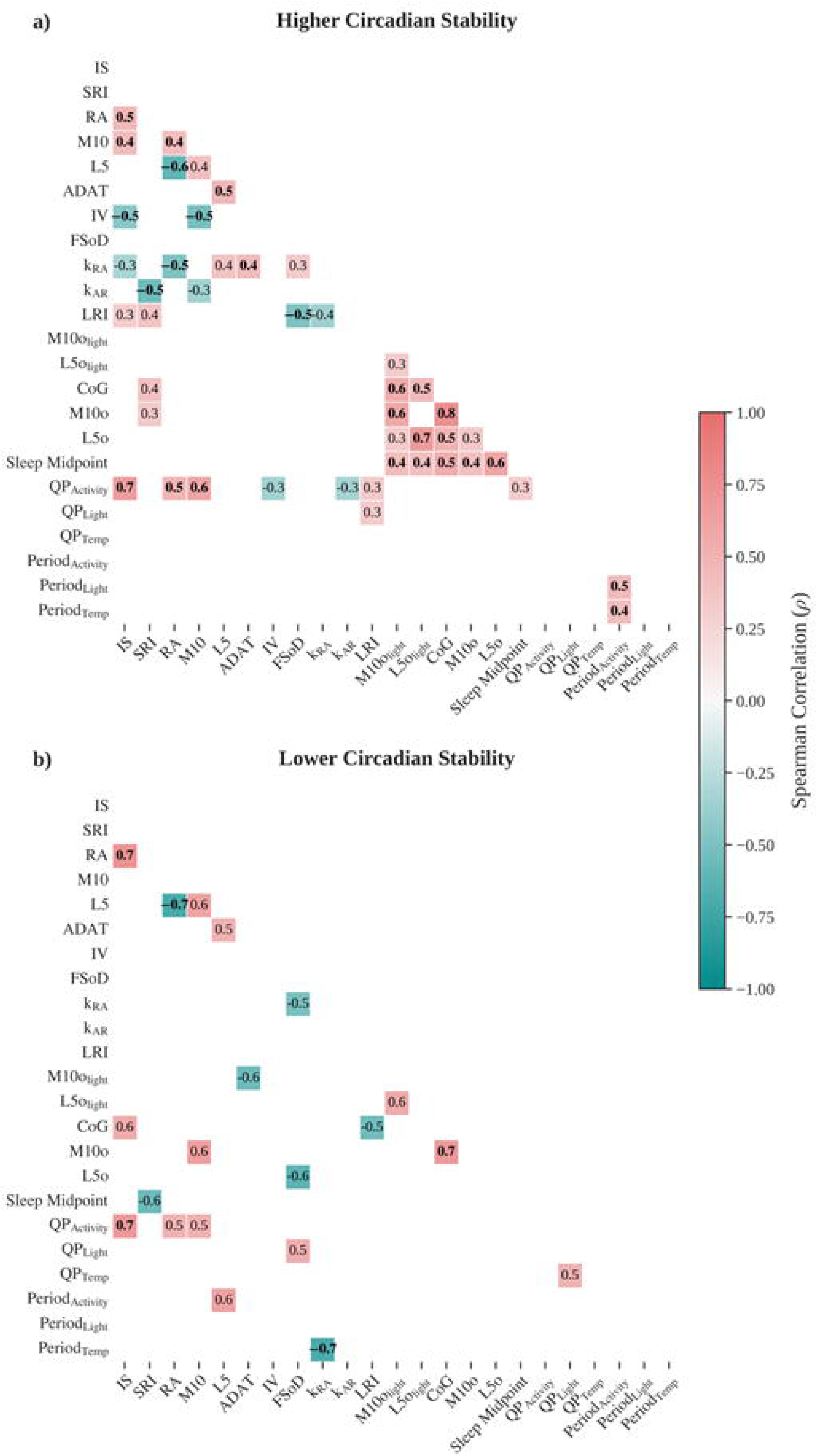
Speannan con-elation matrices for selected rhythm and behavioral variables within each circadian stability cluster. (a) Higher Circadian Stability group. (b) Lower Circadian Stability group. Only significant con-elations are shown (p > 10.31, FDR< 0.05), with bold values indicating **p < 0.01.** Color scale represents the strength and direction of the con-elation (red= positive, blue = negative).

In contrast, the Lower Circadian Stability (LCS) cluster exhibited fewer significant associations. These were limited to relationships between stability metrics and amplitude or rhythm prominence (IS–RA ρ = 0.74; IS–QPActivity ρ = 0.70), a negative association between relative amplitude and activity during the least active 5-h period (RA–L5 ρ = –0.71), an association between behavioral fragmentation and the temperature rhythm period (kAR–PeriodTemperature ρ = –0.69), and a single phase-related association between M10o and CoG (ρ = 0.69) (Figure 3B).

### Predictive Value of Clusters Versus Pupillary Reactivity

To determine the relative predictive power of cluster membership and pupillary reactivity, we conducted linear regressions using each of these factors as independent variables for the key circadian metrics. The results revealed that cluster membership had greater explanatory value than pupillary reactivity across most variables.

As shown in Table 6, cluster membership significantly predicted variation in core metrics such as RA (R² = 0.61), IS (R² = 0.54), and QPActivity (R² = 0.48), all with p-values < 0.001. Pupillary reactivity, by contrast, showed a more modest association with rhythmicity. Among the reactivity-based models (Table 7), only RA and L5 reached statistical significance at the p < 0.001 level, and their effect sizes were consistently lower than those observed for cluster-based regressions.

These findings reinforce the notion that endogenous rhythm organisation, as captured by clustering of multidimensional circadian features, provides a more robust explanation of rest–activity dynamics than pupillary light response alone.

### Pupillary reactivity and light perception by cluster

To investigate whether the distribution of clusters differed according to the combined effects of pupillary reactivity and light perception (Figure 4), a chi-squared test was performed. No significant association was found between the categorical variable Reactivity × Light Perception and cluster membership. To further explore this relationship, a logistic regression model was fitted with *Cluster* as the dependent variable (HCS,LCS), and Pupillary Reactivity, Light Perception, and their interaction as predictors. The model did not reach statistical significance. None of the predictors showed independent associations with cluster membership, namely Reactivity ; Light perception and Interaction term

**Figure 4.**
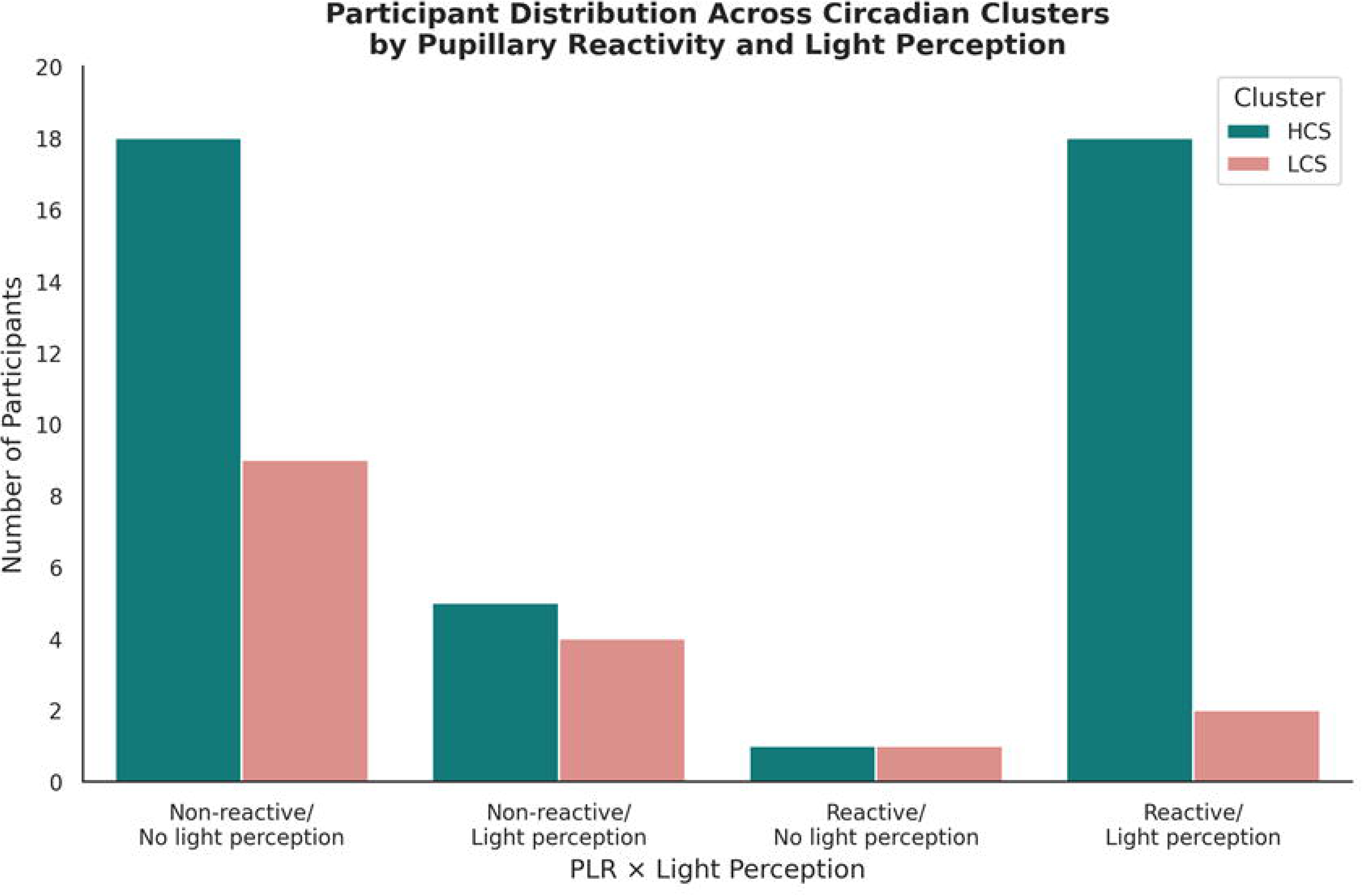
Distribution of pa1ticipants across clusters (HCS and LCS) according to the combination of pupilla1y reactivity and light perception. Bar height indicates the number of individuals in each group.

## Discussion

In this study, we implemented a semi-supervised machine learning analytical pipeline to investigate whether refined multivariate approaches based on non-parametric circadian metrics could detect subtle alterations in circadian rhythmicity beyond those captured by traditional parametric methods. Circadian organisation was first examined using pupillary light reactivity (PLR) as a functional proxy of intrinsically photosensitive retinal ganglion cell (ipRGC) integrity, given their role in conveying photic information to the suprachiasmatic nucleus (SCN) (Markwell et al., 2010; Smith et al., 2020). In this context, rest–activity rhythms were initially characterised using spectral estimates of rhythm period and phase-angle relationships. We then applied a data-driven analytical pipeline combining unsupervised clustering across multiple circadian metrics with SHAP-based feature importance analyses to identify circadian phenotypes and determine the rhythmic features most relevant for cluster separation. Finally, clusters were characterised through between-group comparisons, within-cluster correlations among circadian metrics, and regression models. Together, this analytical strategy enabled an integrated characterisation of the identified circadian phenotypes, capturing the multidimensional nature of circadian organisation.

Spectral analysis revealed entrainment of RAR in all participants with preserved PLR and in 80.6% of non-reactive individuals, indicating that rest–activity rhythms can maintain a 24-h period even in the absence of photic responsiveness. These findings are notable given the well-documented vulnerability of blind individuals to chronodisruption, including Non-24 h sleep disorder. Prior investigations, based on melatonin rhythms analyzed using cosinor methods, reported that fewer than 40% of individuals without light perception are normally entrained (Skene et al., 1999; Lockley et al., 2008; Flynn-Evans et al., 2014). While differences in methodology and analytical approaches may contribute to this discrepancy, actigraphy is widely recognised as a reliable tool for assessing circadian organisation (Leocadio-Miguel & Araújo, 2022), as its measures correlate strongly with circadian phase markers, such as 6-sulphatoxymelatonin (aMT6s) (Lockley et al., 1999), and polysomnography evaluations of sleep–wake disorders (Roh et al., 2022).

However, while spectral analysis provides valuable information regarding periodicity, it does not fully capture the structural complexity and integrity of circadian organisation. To move beyond period estimation, we evaluated multiple non-parametric actigraphy metrics (Hammad et al., 2021) by applying a semi-supervised machine learning approach, revealing two distinct groups determined primarily by: Relative Amplitude (RA), Interdaily Stability (IS), and rhythm strength (QPActivity). Given that these metrics collectively reflect the stability and robustness of the rest–activity cycle, we designated the resulting circadian phenotypes as HCS and LCS. Although only seven participants were classified as non-entrained based on spectral criteria, the clustering approach identified nine additional individuals who, despite exhibiting a 24-hour rest–activity rhythm, demonstrated multidimensional patterns consistent with reduced structural stability. Consequently, the LCS group comprised 28% of the sample, including three reactive and thirteen non-reactive participants. As a result, 72% of the total sample were classified within the HCS phenotype, which still represents one of the highest proportions of consolidated circadian organisation reported in blind adults to date.

Notably, the magnitude of stability-related indices observed in the HCS phenotype closely paralleled values reported in large community-based adult samples. Mitchell et al. (2017) provided a well-characterised population reference by examining rest–activity rhythm distributions in 590 free-living adults across multiple U.S. metropolitan regions, using seven consecutive days of wrist actigraphy collected under ecologically valid, non-interventional conditions. In that setting, non-parametric circadian indices varied systematically across lifespan and demographic strata, reinforcing their validity as markers of real-world circadian organisation rather than laboratory-derived constructs. Mean (±SD) values in that cohort were RA = 0.77 ± 0.11, IS = 0.49 ± 0.12, and IV = 0.86 ± 0.24. In comparison, our HCS group demonstrated RA = 0.79 ± 0.08, IS = 0.49 ± 0.10, and IV = 0.79 ± 0.18, indicating substantial quantitative overlap with population benchmarks.

Large-scale accelerometry data from the UK Biobank further contextualise these findings. In more than 20,000 middle-aged adults, mean values for intra-daily variability ranged from 0.6 to 0.7, inter-daily stability from 0.6 to 0.7, and relative amplitude around 0.85, with Sleep Regularity Index values between 58 and 63 (Walker et al., 2024). Although these values are modestly higher than those observed in our cohort, they derive from an occupationally active and socioeconomically structured sample, in which daily routines are likely strongly shaped by employment demands and social timing. Taken together, these comparisons suggest that circadian organisation in blindness is not categorically distinct from that observed in sighted free-living populations, but instead occupies a range within the broader normative distribution. This appears to be at least correct for the blind population living at low latitudes.

Among the potential explanations for this phenomenon, it is important to consider that many foundational studies using objective metrics of rhythmicity in blind populations were conducted decades ago. Today, technological advancements, cultural habits, and modern social dynamics may contribute to our findings by introducing novel or intensified non-photic cues (Khan & Khusro, 2021). However, the extent to which these factors influence circadian organisation in blind individuals remains unclear and warrants further investigation in future studies.

Nevertheless, a classic study by Klerman et al. (1998), which elegantly integrated real-world and laboratory data, demonstrated that 9 out of 15 totally blind individuals (60%) living in society remained naturally entrained to the 24-hour cycle. Their study, conducted with healthy participants ranging from 23 to 70 years old, showed that scheduled sleep–wake cycles under near darkness (<0.03 lx), combined with a brief 10-minute bout of stationary cycling, were sufficient to induce phase shifts in circadian rhythms. Thus, the authors argue that non-photic zeitgebers, such as structured physical activity and rigid behavioral routines, can exert a significant synchronizing effect on the human circadian pacemaker, successfully compensating for the absence of photic input.

For such behavioural routines to sustain long-term circadian alignment without light, a consistent environmental scenario is paramount. Therefore, an important explanatory factor in our study is the ecological context of our cohort. This study was conducted at 5°S, a tropical latitude characterised by minimal seasonal variation in photoperiod and relatively stable thermal conditions throughout the year. In sighted individuals, latitude has been seen as an important factor in modulating chronotype and circadian rhythms. Our group demonstrated in a cohort of 12,884 volunteers living in the same time zone, with different latitudes, Natal (5°46′ south, 35°12′ west), São Paulo (23°33′ south, 46°37′ west), and Porto Alegre (30°2′ south, 51°13′ West), morningness was associated with low latitudes and eveningness with higher latitudes (Leocadio-Miguel et.al., 2017).

Recent studies conducted in sighted populations consistently report a relationship between latitude and sleep–wake patterns, suggesting that geographic context is a universal modulator of human timing. Large-scale data from over 400,000 U.S. adults demonstrate that geographic context, specifically latitude and time zone, plays a more consistent role than seasonality in shaping sleep duration (DelRosso & Vodapally, 2026). These findings align with Chilean population studies spanning a broad latitudinal range, which similarly show longer sleep at higher latitudes, particularly on weekends (Brockmann et al., 2017). Furthermore, year-long objective monitoring in 4,683 U.S. physicians recently refined this association, showing that each 1° increase in latitude correlates with longer total sleep time. While simulation analyses in that cohort demonstrated seasonally specific photoperiod-mediated effects, residual direct effects of latitude persisted (Ross et al., 2025), reinforcing the notion that geographic location imposes a stable and continuous influence on human circadian organisation.

These studies acknowledge that while photoperiod is the primary pathway linking latitude to sleep, it does not fully account for geographic variation. Factors such as temperature and solar radiation are tightly coupled to the light–dark cycle in equatorial regions like ours. Temperature is known to influence circadian rhythmicity at both central and peripheral levels. Two primary models have been proposed to explain temperature-based modulation of the circadian system: the network model, which attributes thermal resistance to neuronal coupling within the suprachiasmatic nucleus (SCN) (Buhr et al., 2010), and the pathway model, which suggests a dedicated thermosensory signaling route to the clock (Kidd et al., 2015). Furthermore, peripheral clocks respond to thermal cues through mechanisms such as alternative splicing (e.g., in *U2af26*) and the activation of heat shock pathways. These findings raise the intriguing possibility that, in the absence of photic responsiveness, consistent thermal inputs combined with regular behavioral organisation may support circadian alignment in visually impaired individuals (Ball et al., 2025).

Based on this evidence, we hypothesise that in regions with pronounced seasonal variations in photoperiod and temperature, blind individuals are exposed to greater environmental temporal variability, potentially increasing the risk of internal–external misalignment. In contrast, low-latitude settings reduce the demand for repeated seasonal re-entrainment, offering a highly consistent temporal scenario wherein behavioral and social zeitgebers can exert maximal stabilizing effects. Importantly, this interpretation does not deny the role of photic mechanisms but instead refines the concept of latitude. Rather than being solely a proxy for seasonal amplitude, it represents a broader ecological temporal regime encompassing solar regularity, thermal stability, and social organisation.

To further validate these estimates, we observed internal coherence within the HCS group and characterised the HCS phenotype.. The behavioural and environmental variables interacted closely, each contributing to rhythm maintenance (Gall & Shuboni-Mulligan, 2022). The center of gravity (CoG) was significantly associated with multiple phase indicators (including M10light, L5light, sleep midpoint, and activity/rest onsets) indicating a coordinated phase orientation across behavioral and environmental domains. Core temporal markers (M10o, L5o, M10light, and L5light) were intercorrelated, demonstrating tight coupling between motor activity and environmental light exposure. Stability-related metrics were similarly integrated: higher IS was associated with lower Intradaily Variability (IV), reduced rest–activity transitions (kRA), greater RA, and higher M10 activity, collectively reflecting the coordinated regulation of rhythm strength and consolidation.

In contrast, the LCS group exhibited few significant correlations between stability and amplitude metrics (e.g., IS–RA and RA–L5), and phase-marker associations were largely restricted to M10 onset and the center of gravity (CoG). Lower relative amplitude (RA) accompanied by increased activity during the least active 5-hour period (L5) is commonly observed when sleep becomes fragmented, as nocturnal awakenings and reduced sleep efficiency increase motor activity during the biological night (Van Someren et al., 1999; Gonçalves et al., 2015).

This pattern was also consistent with the observed values of stability metrics, including a low Sleep Regularity Index (SRI = 38.78 ± 21.82) and low interdaily stability (IS = 0.26 [0.19–0.29]). More broadly, recent large-scale actigraphy studies indicate that sleep regularity is a stronger predictor of mortality risk than sleep duration, and that irregular sleep is associated with higher risks of all-cause, cardiometabolic, and cancer mortality (Windred et al., 2024). Overall, the reduced coherence across circadian features suggests a less organised circadian behavioural structure.

A recent review by Boivin et al. (2022) described how circadian disruption in shift workers involves not only misalignment between behavioral rhythms and the environmental light–dark cycle but also a state of internal desynchronisation across multiple levels of the circadian system. Under such conditions, circadian disruption has been associated with disturbances in sleep and alertness, impaired cognitive performance, increased cardiometabolic risk, metabolic syndrome, type 2 diabetes, cardiovascular disease, gastrointestinal complaints, menstrual irregularities, reproductive difficulties, and elevated risks for certain cancers.

In blind individuals specifically, disruption of the phase relationship between the sleep–wake cycle and the circadian pacemaker has been shown to impair alertness, mood, and daytime functioning, particularly when wakefulness occurs during the biological night. Lockley et al. (2008) demonstrated that disruption of this phase relationship in non-entrained blind individuals resulted in impaired daytime functioning equivalent to that typically experienced when individuals remain awake during the night.

Taken together, these findings support the interpretation that the LCS phenotype may represent a vulnerable circadian behavioral profile characterised by reduced temporal coherence of daily rhythms, with potential relevance for long-term health outcomes. Identifying this phenotype may help recognise individuals with greater circadian instability who could represent a target population for chronobiotic interventions. Nevertheless, further characterisation of this phenotype is needed, particularly to clarify the mechanisms underlying this circadian profile and the factors contributing to its emergence.

Regression analyses further indicated that circadian phenotype classification explained substantially more variance in rhythmic parameters than pupillary light reactivity (PLR). Importantly, these differences remained significant even after adjusting for age and sex in our multivariate models. Neither PLR, nor subjective light perception, significantly predicted phenotype allocation, suggesting that behavioural and environmental integration heavily outweigh residual photic responsiveness in determining circadian stability.

This framework conceptualises circadian rhythms as complex phenotypes emerging from the interaction of stability, amplitude, timing, fragmentation, and environmental exposure. Because it relies on wearable monitoring and computationally reproducible methods, it can be readily extended to populations exposed to circadian challenges, including shift workers, individuals living at higher latitudes, or patients with circadian rhythm sleep–wake disorders. Such applications may help identify vulnerability profiles and inform personalised chronobiological interventions.

However, our results warrant validation in independent cohorts. Future research should prioritise cross-latitudinal comparisons involving blind individuals to precisely identify potential behavioural ‘rhythm stabilisers’. Such investigations are crucial for designing targeted non-photic synchronisation interventions that extend beyond standard pharmacological approaches like exogenous melatonin administration (Leonard-Hawkhead et.al, 2025).

Collectively, our findings support a model in which circadian stability in blindness emerges from the dynamic interaction between photoreceptive capacity, behavioral organisation, and environmental temporal structure. In low-latitude settings characterised by minimal seasonal variability, this interaction may favor the maintenance of consolidated and structurally integrated circadian rhythms. These results refine the interpretation of circadian vulnerability in blindness and underscore the importance of ecological context in shaping temporal organisation.

### Limitations

This study has limitations that should be acknowledged. First, while actigraphy provides valuable estimates of rest–activity rhythms in ecological settings, it reflects behavioural patterns rather than direct biological markers of circadian phase, such as dim light melatonin onset (DLMO) (Lockley et al., 1997). Second, although pupillary light reactivity serves as an accessible proxy for residual non-visual photoreception (Smith et al., 2019), it does not capture the full functional status of the circadian pacemaker. Pupillary reactivity, while indicative of preserved ipRGC function, does not establish the capacity for circadian photoentrainment or melatonin suppression, which require direct physiological assessment (Hull, Czeisler & Lockley, 2018). Third, although demographic variables such as age and sex were included, the potential influence of duration of blindness on circadian outcomes was not systematically explored and warrants further investigation. Nevertheless, the lengthy monitoring periods, the sizable cohort living under environmental stability, and the use of unsupervised clustering to identify emergent phenotypes strengthen the robustness and ecological validity of our findings. Future studies should integrate actigraphy with hormonal and molecular biomarkers, detailed lifestyle tracking, and longitudinal designs to better understand the mechanisms supporting circadian alignment in sensory-deprived populations.

### Conclusion

This study demonstrates that circadian organisation in blind individuals may remain stable even in the absence of pupillary light reflex, particularly under conditions of environmental and behavioural regularity. The high prevalence of entrained rest–activity rhythms observed in this cohort suggests that circadian stability emerges from the interaction between physiological capacity and ecological context, rather than from photic responsiveness alone. Our findings further indicate that multidimensional circadian phenotyping provides a more informative framework than single physiological markers for understanding rhythmic organisation in visually impaired populations. Together, these results support the hypothesis that consistent environmental cues and structured behavioural routines may facilitate compensatory mechanisms that sustain circadian alignment, highlighting the potential importance of ecological context for both research and intervention strategies targeting circadian disturbances in blindness.

## Acknowledgments

In memory of Prof. Malcolm von Schantz, who dedicated his career to the study of chronobiology and inspired a generation of researchers, including several authors of this manuscript. He would, as usual, have had something characteristically sharp to say about it.

We thank all participants for their time and commitment to this study. We are grateful to the team at the Institute of Education and Rehabilitation of the Blind of Natal (IERC) for their collaboration in participant recruitment. We also acknowledge the Graduate Program in Psychobiology at the Federal University of Rio Grande do Norte (UFRN) for institutional support.

This work was supported by the Brazilian National Council for Scientific and Technological Development (CNPq; grants 314197/2021-4 and 406943/2023-0) and by the Coordination for the Improvement of Higher Education Personnel (CAPES), which provided doctoral funding, including support through the Programa de Doutorado Sanduíche no Exterior (PDSE; CAPES Notice No. 06/2024). We also thank Northumbria University for hosting the international doctoral fellowship period, during which part of the data processing and analysis was conducted.

Generative artificial intelligence (AI) was used as a collaborative tool to assist with structural optimisation of the data analysis scripts and English language refinement. All AI content was carefully reviewed and validated by the author, who assumes full responsibility for the integrity of the work.

## Glossary Of Terms

RHT: Retinohypothalamic Tract System;
SCN: Suprachiasmatic Nucleus;
RAR: Rest–Activity Rhythms;
PLR: Pupillary Light Reflex;
HCS: Higher Circadian Stability;
LCS: Higher Circadian Stability

## Reference

1. Allen, A. E. (2019). Circadian rhythms in the blind. Current Opinion in Behavioral Sciences, 30, 73–79. 10.1016/j.cobeha.2019.06.003

2. Ball, D. M., Mann, S. S., Santhi, N., Speekenbrink, M., & Walsh, V. (2025). Temperature as a circadian timing cue in the visually impaired. Progress in Brain Research, 292, 1–24. 10.1016/bs.pbr.2025.02.004

3. Benjamini, Y. & Hochberg, Y. Controlling the false discovery rate: a practical and powerful approach to multiple testing. J. R. Stat. Soc.: Ser. B (Methodol.) 57, 289–300 (1995).

4. Boivin, D. B., Boudreau, P., & Kosmadopoulos, A. (2022). Disturbance of the circadian system in shift work and its health impact. Journal of Biological Rhythms, 37(1), 3–28. 10.1177/07487304211064218

5. Bonmati-Carrion, M. A., et al. (2016). Relationship between human pupillary light reflex and circadian system status. PLOS ONE, 11(9), e0162476. 10.1371/journal.pone.0162476

6. Boudebesse, C., Geoffroy, P. A., Bellivier, F., Henry, C., Folkard, S., Leboyer, M., & Etain, B. (2014). Correlations between objective and subjective sleep and circadian markers in remitted patients with bipolar disorder. Chronobiology International, 31(5), 698–704. 10.3109/07420528.2014.895742

7. Brockmann, P. E., Gozal, D., Villarroel, L., Damiani, F., Nuñez, F., & Cajochen, C. (2017). Geographic latitude and sleep duration: A population-based survey from the Tropic of Capricorn to the Antarctic Circle. Chronobiology international, 34(3), 373–381. 10.1080/07420528.2016.1277735

8. Buchman, A. S., Boyle, P. A., Yu, L., Shah, R. C., Wilson, R. S., & Bennett, D. A. (2012). Total daily physical activity and the risk of AD and cognitive decline in older adults. Neurology, 78(17), 1323–1329. 10.1212/WNL.0b013e3182535d35

9. Buhr, E. D., Yoo, S.-H., & Takahashi, J. S. (2010). Temperature as a universal resetting cue for mammalian circadian oscillators. Science, 330(6002), 379–385. 10.1126/science.1195262

10. Cipolla-Neto, J., & Amaral, F. G. do. (2018). Melatonin as a Hormone: New Physiological and Clinical Insights. Endocrine Reviews, 39(6), 990–1028. 10.1210/er.2018-00084

11. Czeisler, C. A., Shanahan, T. L., Klerman, E. B., Martens, H., Brotman, D. J., Emens, J. S., Klein, T., & Rizzo, J. F. (1995). Suppression of Melatonin Secretion in Some Blind Patients by Exposure to Bright Light. New England Journal of Medicine, 332(1), 6–11. 10.1056/nejm199501053320102

12. David, M. C. M. M., Mattos, M. S. B., Souto, J. J. S., Brito, S. A. C. F., Leite, E. S., Valença, E. N., Galdino, G. S., Sampaio, P. G. G., Moura, D. M. C., Miguel, M. A. L., Araújo, J. F., Franco, C. I. F., & Matos, R. J. B. (2022). Changes in the rest-activity rhythm in migraine patients are associated with anxiety symptoms. Brazilian Journal of Psychiatry. 10.47626/1516-4446-2021-2367

13. Danilevicz, I. M., et al. (2024). Reliable measures of rest–activity rhythm fragmentation: How many days are needed? European Review of Aging and Physical Activity, 21, 29. 10.1186/s11556-024-00364-5

14. Depieri, N.B. (2021) Caracterização do ritmo de atividade e repouso, de exposição à luz e qualidade de sono de sujeitos cegos. Dissertação (Mestrado). 111f. Programa de pós graduação em Psicobiologia, Universidade Federal do Rio Grande do Norte, Natal, Rio Grande do Norte, Brasil

15. Diez-Noguera, A. (1994). A functional model of the circadian system based on the degree of intercommunication in a complex system. *American Journal of Physiology-Regulatory*, Integrative and Comparative Physiology, 267(4), R1118–R1135. 10.1152/ajpregu.1994.267.4.r1118

16. Emens, J. S., Laurie, A. L., Songer, J. B., & Lewy, A. J. (2013). Non-24-Hour Disorder in Blind Individuals Revisited: Variability and the Influence of Environmental Time Cues. SLEEP, 36(7), 1091–1100. 10.5665/sleep.2818

17. Flynn-Evans, E. E., et al. (2014). Circadian rhythm disorders and melatonin production in 127 blind women with and without light perception. Journal of Biological Rhythms, 29(3), 215–224. 10.1177/0748730414536852

18. Flynn-Evans, E. E., & Lockley, S. W. (2016). A Pre-Screening Questionnaire to Predict Non-24-Hour Sleep-Wake Rhythm Disorder (N24HSWD) among the Blind. Journal of Clinical Sleep Medicine, 12(05), 703–710. 10.5664/jcsm.5800

19. Forner-Cordero, A., Umemura, G. S., Furtado, F., & Gonçalves, B. da S. B. (2018). Comparison of sleep quality assessed by actigraphy and questionnaires to healthy subjects. Sleep Science, 11(3), 141–145. 10.5935/1984-0063.20180027

20. Forbes, E. E., Dahl, R. E., Almeida, J., Ferrell, R. E., Nimgaonkar, V. L., Mansour, H., Sciarrillo, S. R., Holm, S. M., Rodriguez, E. E., & Phillips, M. L. (2012). PER2 rs2304672 Polymorphism Moderates Circadian-Relevant Reward Circuitry Activity in Adolescents. 71(5), 451–457. 10.1016/j.biopsych.2011.10.012

21. Gall, A. J., & Shuboni-Mulligan, D. D. (2022). Keep Your Mask On: The Benefits of Masking for Behavior and the Contributions of Aging and Disease on Dysfunctional Masking Pathways. Frontiers in Neuroscience, 16. 10.3389/fnins.2022.911153

22. Gonçalves, B., Adamowicz, T., Louzada, F. M., Moreno, C. R., & Araujo, J. F. (2015). A fresh look at the use of nonparametric analysis in actimetry. Sleep Medicine Reviews, 20, 84–91. 10.1016/j.smrv.2014.06.002

23. Hammad, G. B., Reyt, M., Beliy, N., Baillet, M., Deantoni, M., Lesoinne, A., Muto, V., & Schmidt, C. (2021). pyActigraphy: Open-source python package for actigraphy data visualization and analysis. 17(10), e1009514–e1009514. 10.1371/journal.pcbi.1009514

24. Hammad, G., Wulff, K., Skene, D. J., Mirjam Münch, & Spitschan, M. (2024). Open-Source Python Module for the Analysis of Personalized Light Exposure Data from Wearable Light Loggers and Dosimeters. Leukos (Online), 1–10. 10.1080/15502724.2023.2296863

25. Hand, A. J., Stone, J. E., Shen, L., Vetter, C., Cain, S. W., Bei, B., & Phillips, A. J. K. (2023). Measuring Light Regularity: Sleep Regularity is Associated with Regularity of Light Exposure in Adolescents. Sleep. 10.1093/sleep/zsad001

26. Hull, J. T., Czeisler, C. A., & Lockley, S. W. (2018). Suppression of Melatonin Secretion in Totally Visually Blind People by Ocular Exposure to White Light. Ophthalmology, 125(8), 1160–1171. 10.1016/j.ophtha.2018.01.036

27. Hut, R. A., Paolucci, S., Dor, R., Kyriacou, C. P., & Daan, S. (2013). Latitudinal clines: an evolutionary view on biological rhythms,. Proceedings of the Royal Society B: Biological Sciences, 280(1765), 20130433. 10.1098/rspb.2013.0433

28. Jolliffe, I. T., & Cadima, J. (2016). Principal component analysis: a review and recent developments. *Philosophical Transactions of the Royal Society A: Mathematical*, Physical and Engineering Sciences, 374(2065), 20150202. 10.1098/rsta.2015.0202

29. Khan, A., & Khusro, S. (2021). An insight into smartphone-based assistive solutions for visually impaired and blind people: Issues, challenges and opportunities. Universal Access in the Information Society, 20, 265–298. 10.1007/s10209-020-00733-8

30. Kenagy, G. J. (1980). Center-of-gravity of circadian activity and its relation to free-running period in two rodent species. Journal of Interdisciplinary Cycle Research, 11(1), 1–8. 10.1080/09291018009359682

31. Kidd, P. B., et al. (2015). Temperature compensation and temperature sensation in the circadian clock. PNAS, 112(46), E6284–E6292. 10.1073/pnas.1511215112

32. Klerman, E. B., Rimmer, D. W., Dijk, D.-J., Kronauer, R. E., Rizzo, J. F., & Czeisler, C. A. (1998). Nonphotic entrainment of the human circadian pacemaker. American Journal of Physiology-Regulatory, Integrative and Comparative Physiology, 274(4), R991–R996. 10.1152/ajpregu.1998.274.4.r991

33. Lee, S. Y., & Mesfin, F. B. (2020). Blindness. In StatPearls. StatPearls Publishing. https://www.ncbi.nlm.nih.gov/books/NBK448182/

34. Leonard-Hawkhead, B., Piyasena, M. P., Peto, T., Virgili, G., van Nispen, R. M. A., & Curran, K. (2025). Sleep improvement strategies for people with vision impairment: a scoping review. BMJ Open, 15(12), e113100. 10.1136/bmjopen-2025-113100

35. Leocadio-Miguel, M.A., Fontenele-Araújo, J. (2022). Actigraphy. In: Frange, C., Coelho, F.M.S. (eds) Sleep Medicine and Physical Therapy. Springer, Cham. 10.1007/978-3-030-85074-6_37

36. Leocadio-Miguel, M. A., Louzada, F. M., Duarte, L. L., Areas, R. P., Alam, M., Freire, M. V., Fontenele-Araujo, J., Menna-Barreto, L., & Pedrazzoli, M. (2017). Latitudinal cline of chronotype. Scientific Reports, 7(1). 10.1038/s41598-017-05797-w

37. Lockley, S. W., Skene, D. J., Arendt, J., Tabandeh, H., Bird, A. C., & Defrance, R. (1997). Relationship between Melatonin Rhythms and Visual Loss in the Blind1. The Journal of Clinical Endocrinology & Metabolism, 82(11), 3763–3770. 10.1210/jcem.82.11.4355

38. Lockley, S. W., Skene, D. J., Butler, L. J., & Arendt, J. (1999). Sleep and Activity Rhythms are Related to Circadian Phase in the Blind. Sleep, 22(5), 616–623. 10.1093/sleep/22.5.616 Lockley, S. W., Arendt, J., & Skene, D. J. (2007). Visual impairment and circadian rhythm disorders. Dialogues in 10.31887/DCNS.2007.9.3/slockley

39. Lockley, S. W., Dijk, D.-J., Kosti, O., Skene, D. J., & Arendt, J. (2008). Alertness, mood and performance rhythm disturbances associated with circadian sleep disorders in the blind. Journal of Sleep Research, 17(2), 207–216. 10.1111/j.1365-2869.2008.00656.x

40. MSD Manual Professional Edition. (n.d.). Assessing visual acuity. Retrieved March 16, 2026, from https://www.msdmanuals.com/professional/multimedia/table/assessing-visual-acuity

41. Markwell, E. L., Feigl, B., & Zele, A. J. (2010). Intrinsically photosensitive melanopsin retinal ganglion cell contributions to the pupillary light reflex and circadian rhythm. Clinical and Experimental Optometry, 93(3), 137–149. 10.1111/j.1444-0938.2010.00479.x

42. Mitchell, J. A., Quante, M., Godbole, S., James, P., Hipp, J. A., Marinac, C. R., Mariani, S., Cespedes Feliciano, E. M., Glanz, K., Laden, F., Wang, R., Weng, J., Redline, S., & Kerr, J. (2017). Variation in actigraphy-estimated rest-activity patterns by demographic factors. Chronobiology international, 34(8), 1042–1056. 10.1080/07420528.2017.1337032

43. Monsivais-Velazquez, D., Bhattacharya, K., Ghosh, A., Dunbar, R. I. M., & Kaski, K. (2017). Seasonal and geographical impact on human resting periods. Scientific Reports, 7(1), 1–10. Article 10717. 10.1038/s41598-017-11125-z

44. Ponce-Bobadilla, A. V., Schmitt, V., Maier, C. S., Mensing, S., & Stodtmann, S. (2024). Practical guide to SHAP analysis: Explaining supervised machine learning model predictions in drug development. Clinical and Translational Science, 17(11). 10.1111/cts.70056

45. Randler, C. (2008). Morningness-Eveningness Comparison in Adolescents from Different Countries around the World. Chronobiology International, 25(6), 1017–1028. 10.1080/07420520802551519

46. Randler, C., & Rahafar, A. (2017). Latitude affects Morningness-Eveningness: evidence for the environment hypothesis based on a systematic review. Scientific Reports, 7(1). 10.1038/srep39976

47. Refinetti R, Cornélissen G, Halberg F. Procedures for numerical analysis of circadian rhythms. Biol Rhythm Res. 2007;38(4):275–325. doi: 10.1080/09291010600903692. PMID: 23710111; PMCID: PMC3663600.

48. Roh, H. W., Choi, S. J., Jo, H., Kim, D., Choi, J., Son, S. J., & Joo, E. Y. (2022). Associations of actigraphy derived rest activity patterns and circadian phase with clinical symptoms and polysomnographic parameters in chronic insomnia disorders. Scientific Reports, 12(1), 4895. 10.1038/s41598-022-08899-2

49. Ross, K. E., Pereira-Lima, K., Shedden, K., Burmeister, M., & Sen, S. (2025). Associations of latitude and photoperiod with sleep duration in a yearlong study of US physicians. Sleep medicine, 136, 106840. 10.1016/j.sleep.2025.106840

50. Sack, R. L., et al. (1992). Circadian rhythm abnormalities in totally blind people: Incidence and clinical significance. Journal of Clinical Endocrinology & Metabolism, 75(1), 127–134.

51. Sack, R. L., et al. (2000). Entrainment of free-running circadian rhythms by melatonin in blind people. The New England Journal of Medicine, 343(15), 1070–1077. 10.1056/NEJM200010123431503

52. Skene, D. J., Lockley, S. W., & Arendt, J. (1999). Melatonin in Circadian Sleep Disorders in the Blind. Neurosignals, 8(1-2), 90–95. 10.1159/000014575

53. Skene, D. J., & Arendt, J. (2006). Human circadian rhythms: physiological and therapeutic relevance of light and melatonin. Annals of Clinical Biochemistry, 43(5), 344–353. 10.1258/000456306778520142

54. Smith, J., Flower, O., Tracey, A., & Johnson, P. (2020). A comparison of manual pupil examination versus an automated pupillometer in a specialised neurosciences intensive care unit. Australian Critical Care, 33(2), 162–166. 10.1016/j.aucc.2019.04.005

55. Sokolove, P. G., & Bushell, W. N. (1978). The chi square periodogram: Its utility for analysis of circadian rhythms. Journal of Theoretical Biology, 72(1), 131–160. 10.1016/0022-5193(78)90022-x

56. Van Gelder, R. N. (2008). Non-Visual Photoreception: Sensing Light without Sight. Current Biology, 18(1), R38–R39. 10.1016/j.cub.2007.11.027

57. Van Someren, E. J. W., et al. (1999). Bright Light Therapy: Improved Sensitivity to Its Effects on Rest-Activity Rhythms in Alzheimer Patients by Application of Nonparametric Methods. Chronobiology International, 16(4), 505–518. 10.3109/07420529908998724

58. Walker, B., et al. (2024). Cross-sectional relationships of circadian misalignment and rest-activity rhythms with occupational attainment in UK Biobank participants. Chronobiology International, 1–15. 10.1080/07420528.2024.2441192

59. Windred, D. P., Burns, A. C., Lane, J. M., Saxena, R., Rutter, M. K., Cain, S. W., & Phillips, A. J. K. (2024). Sleep regularity is a stronger predictor of mortality risk than sleep duration: A prospective cohort study. Sleep, 47(1), zsad253. 10.1093/sleep/zsad253

60. Witting, W., et al. (1990). Alterations in the circadian rest-activity rhythm in aging and Alzheimer’s disease. Biological Psychiatry, 27(6), 563–572. 10.1016/0006-3223(90)90523-5

61. Wright, K. P., Jr, Hughes, R. J., Kronauer, R. E., Dijk, D. J., & Czeisler, C. A. (2001). Intrinsic near-24-h pacemaker period determines limits of circadian entrainment to a weak synchronizer in humans. Proceedings of the National Academy of Sciences of the United States of America, 98(24), 14027–14032. 10.1073/pnas.201530198

62. Zaidi, F. H., Hull, J. T., Peirson, S. N., Wulff, K., Aeschbach, D., Gooley, J. J., … & Lockley, S. W. (2007). Short-wavelength light sensitivity of circadian, pupillary, and visual awareness in humans lacking an outer retina. Current biology, 17(24), 2122–2128. 10.1016/j.cub.2007.11.034

