## Supplementary figures and images for "Low-latitude environmental regularity sustains non-photic entrainment in blind adults"

### Supplementary Figure 1

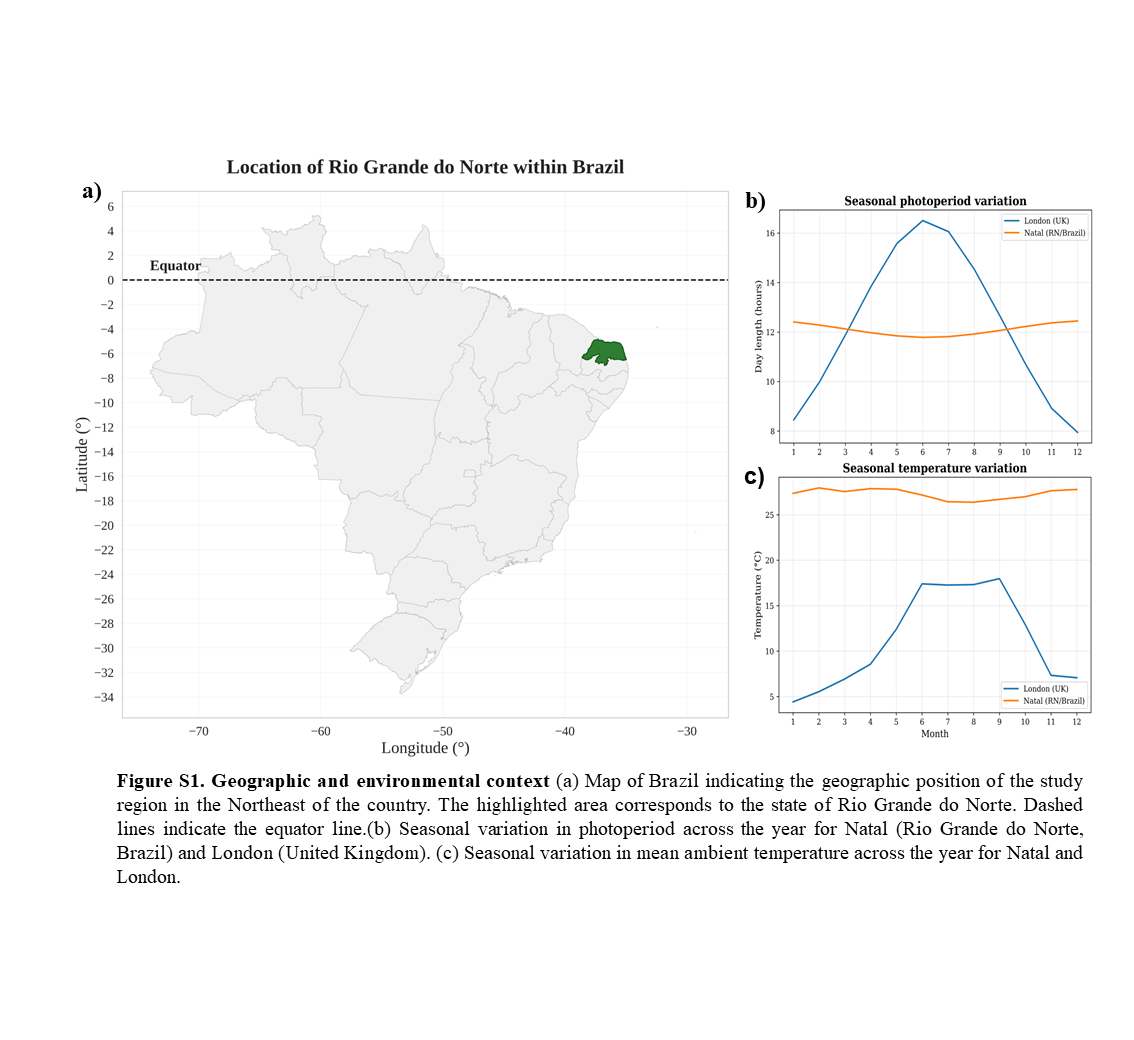

### Supplementary Figure 2

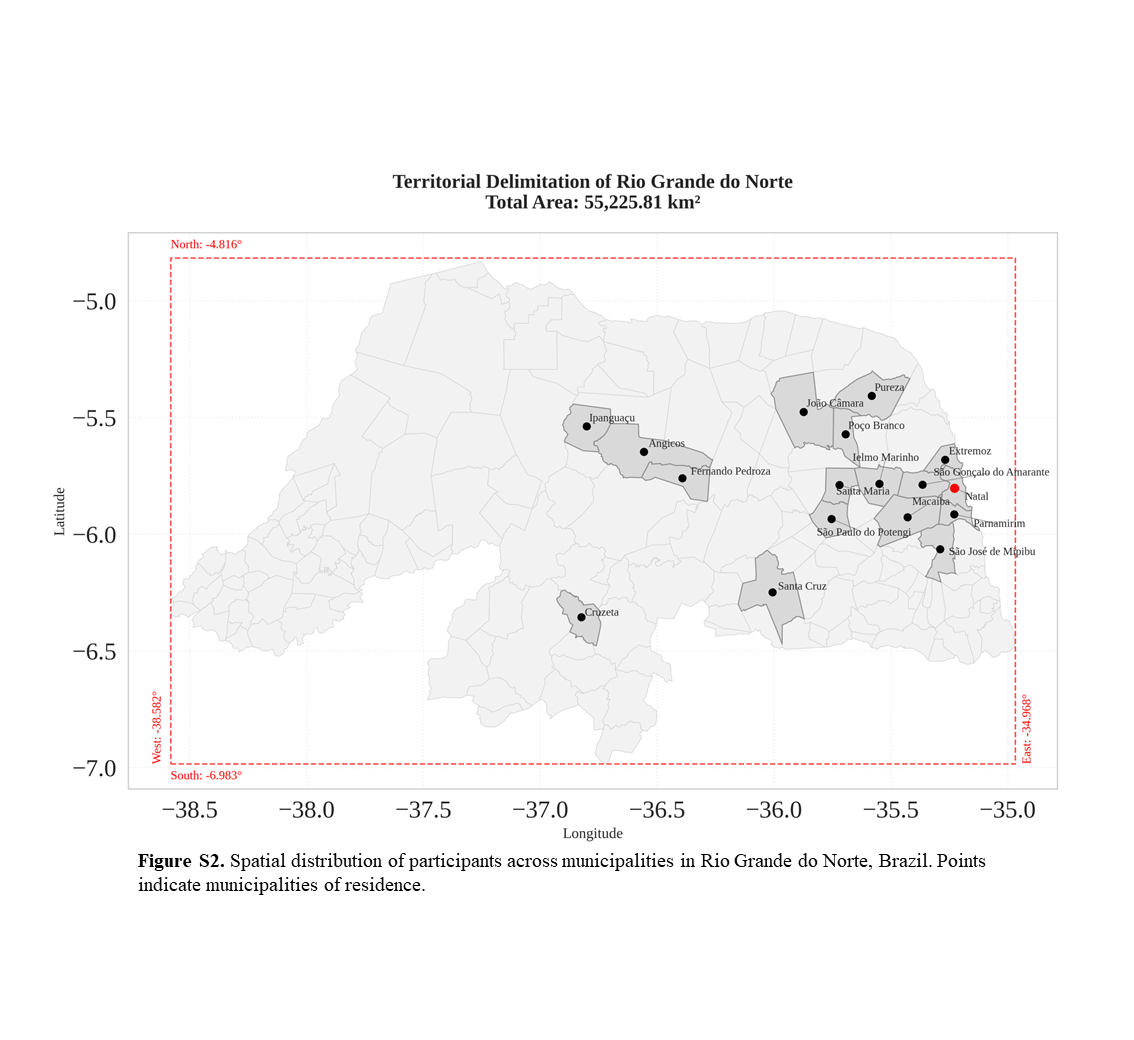

### Supplementary Figure 3

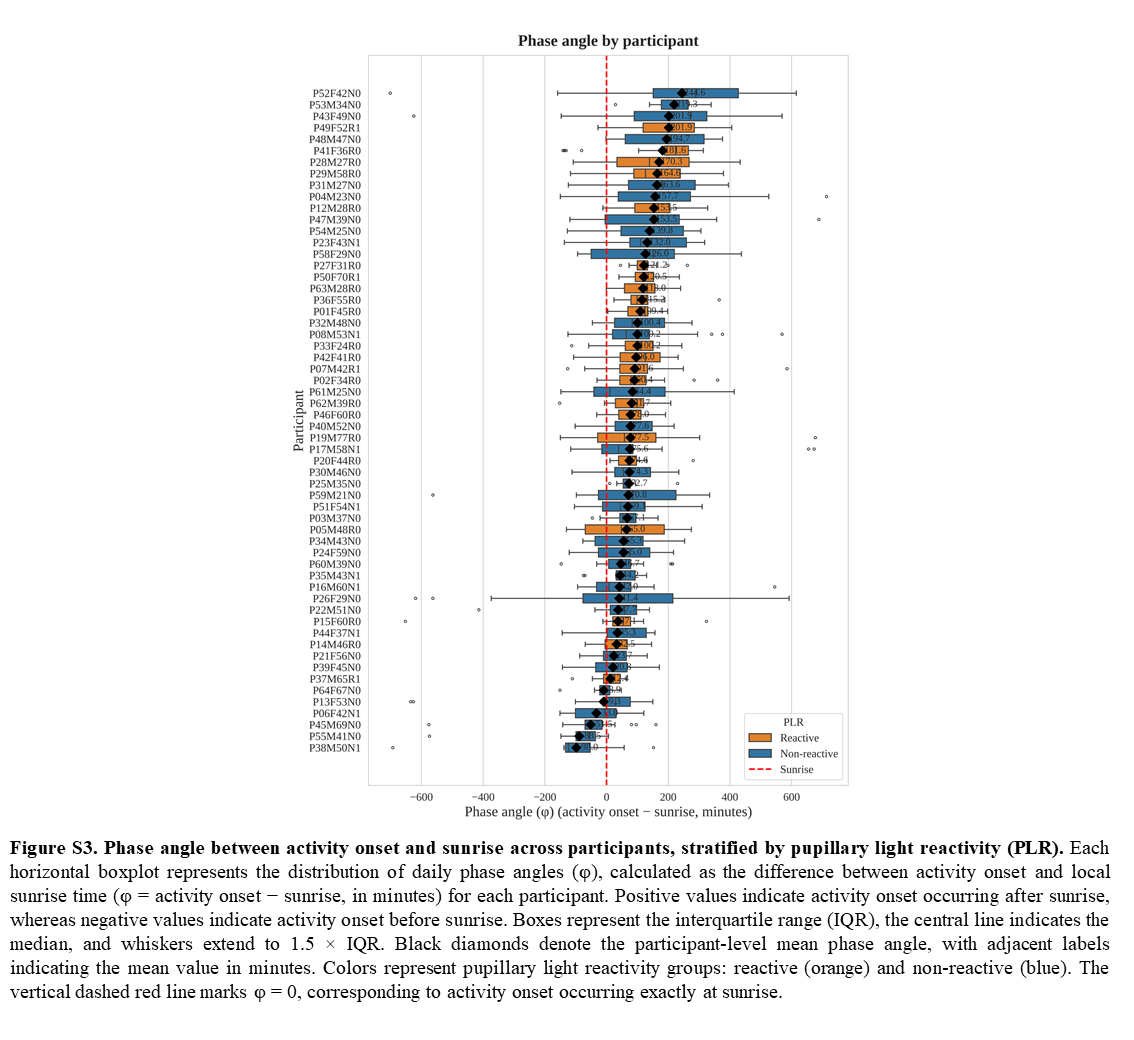

### Supplementary Figure 4

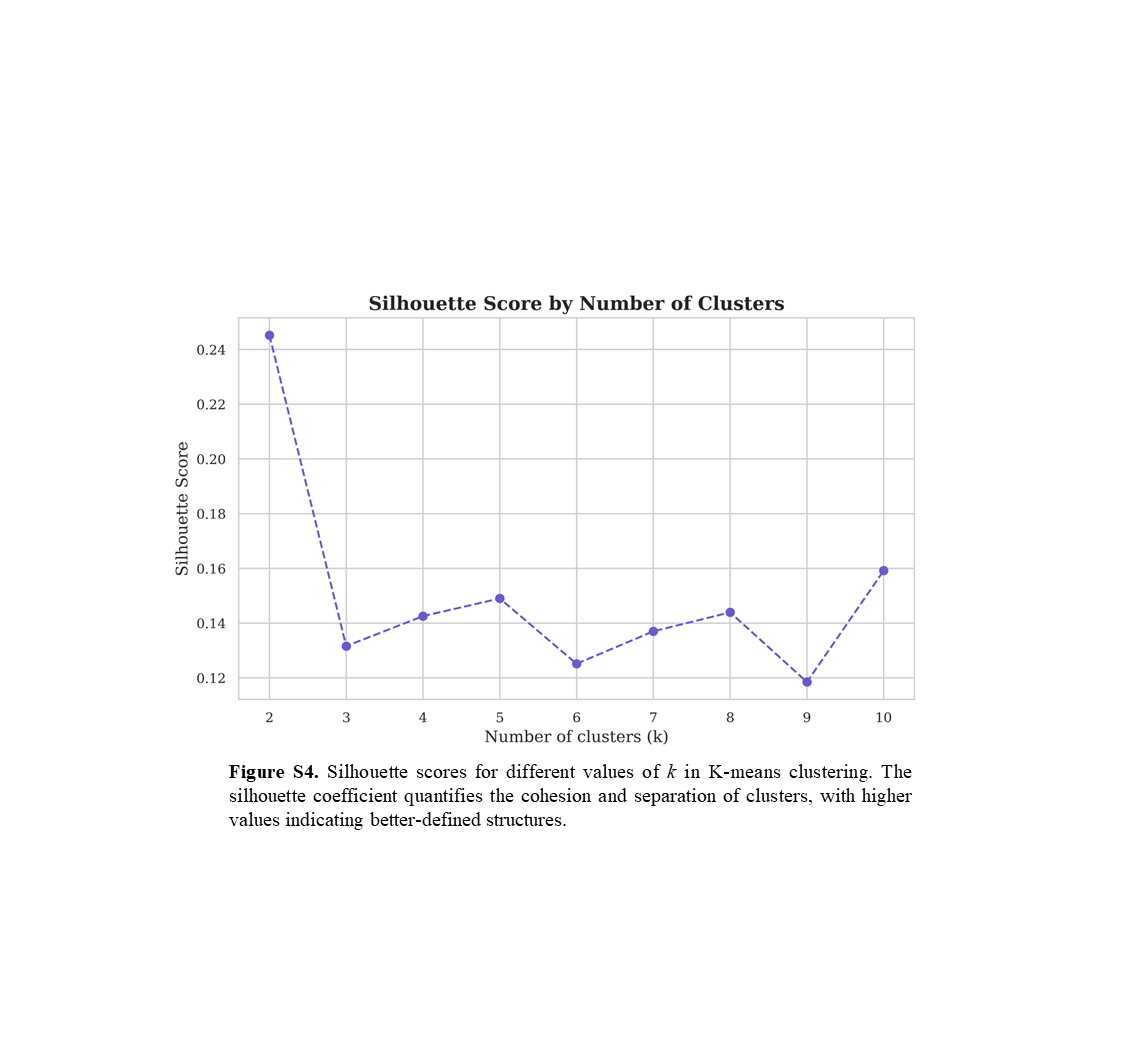

### Supplementary Table 1

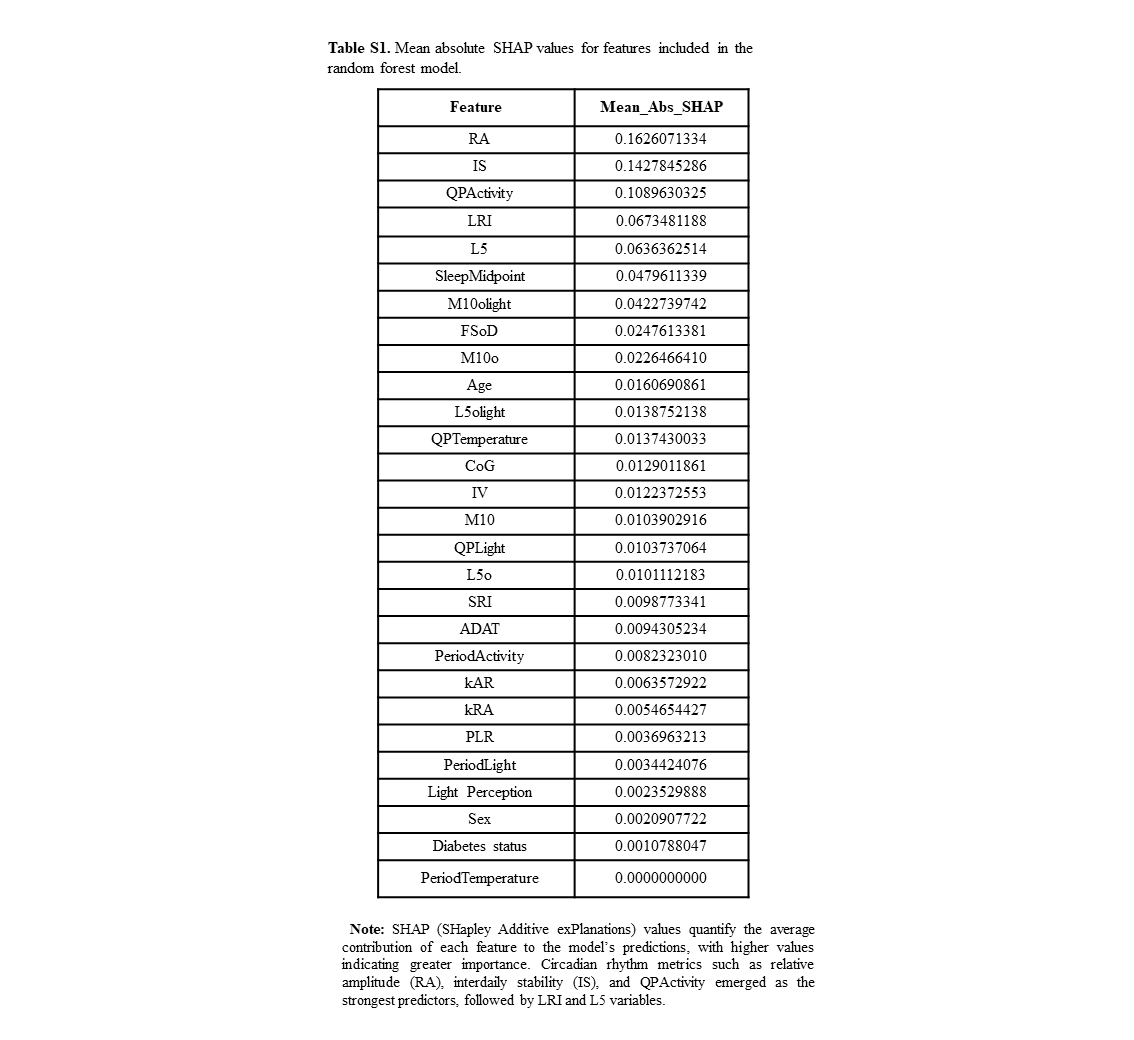

### Supplementary Table 2

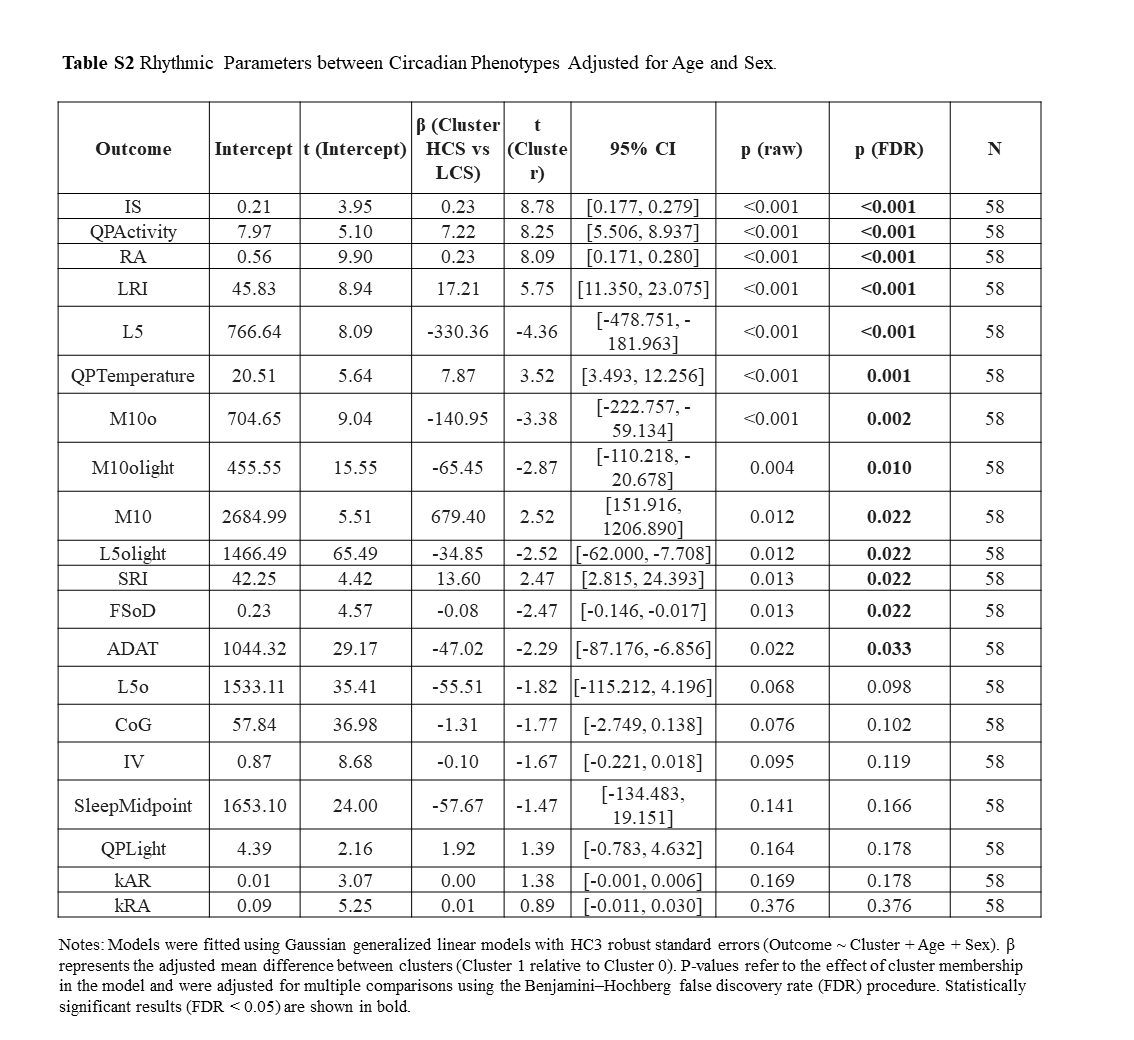
